# Cued unpredictable intermittent access exacerbates loss of control over ethanol drinking

**DOI:** 10.64898/2026.03.31.715677

**Authors:** Eric H Mitten, Jada M. Caldwell, Gerardo Zambrano, Nathaly M Arce Soto, Elizabeth J Glover

## Abstract

**Background:** Loss of control over drinking is a hallmark feature of alcohol use disorder (AUD) that is modeled preclinically through escalation of ethanol consumption and aversion-resistant drinking. Prior work with other reinforcers suggests that within-session unpredictable, intermittent access (uIntA) promotes these phenotypes. However, the effect of uIntA on voluntary ethanol consumption is unknown.

**Methods:** Male and female Long-Evans rats (*n*=9-10/group) underwent seven weeks of daily voluntary ethanol (20% v/v) drinking sessions under either a continuous access (ContA) or uIntA schedule. Following four weeks of baseline, rats were rendered dependent using a two-week chronic intermittent ethanol vapor exposure procedure. Daily testing was maintained through one week into withdrawal from vapor exposure. On the final day of testing, ethanol was adulterated with quinine (30 mg/L) to assess aversion-resistant drinking.

**Results:** Rats drinking under ContA and uIntA exhibited similar levels of average daily ethanol consumption at baseline. However, uIntA elicited a more robust dependence-induced escalation of ethanol consumption compared to ContA, with uIntA sustaining escalation through early withdrawal. Additionally, while rats with ContA to ethanol remained sensitive to quinine even after chronic ethanol vapor exposure, uIntA promoted aversion-resistant drinking in ethanol dependent rats.

**Conclusions:** These results demonstrate that, compared to ContA, uIntA maintains ethanol drinking and exacerbates dependence-induced escalation and aversion-resistant ethanol consumption. This work positions uIntA as a powerful tool to assess psychological and neurobiological factors that may underlie loss of control over drinking.

## Introduction

Alcohol use disorder (AUD) is a chronic, relapsing disorder characterized by several hallmark symptoms, including loss of control over drinking and drinking despite negative consequences (American Psychiatric Association, 2022; Koob, 2024; MacKillop et al., 2022). These symptoms are often investigated preclinically using models of escalated ethanol intake (Becker, 2013; Bowen et al., 2022) and aversion-resistant drinking, respectively (De Oliveira Sergio et al., 2023; Hopf & Lesscher, 2014). These phenotypes are thought to reflect the transition from moderate to intense, problematic intake characteristic of AUD and other substance use disorders (SUD) (Edwards & Koob, 2013). In attempts to more accurately model patterns of human drug intake and seeking to inform the development of SUD-related phenotypes, a bevy of prior work has shown that variable, intermittent timing of reinforcer access can promote uncontrolled intake and responding for both drug and non-drug reinforcers (Allain & Samaha, 2019; Becker, 2013; Clark & Zack, 2023; Spear, 2020). With respect to ethanol, perhaps the most well-characterized effect of schedule on drinking has been observed in the *ad libitum* two bottle choice (2BC) home cage drinking paradigm, which consists of concurrent access to water and ethanol (Bowen et al., 2022; Carnicella et al., 2014; Spear, 2020). Intermittent access 2BC (IA2BC), often defined as limiting 24 hr 2BC sessions to non-consecutive days of the week, can drive significantly higher intake and promote a more robust binge drinking phenotype than continuous 2BC, defined as consecutive 24 hr 2BC sessions (Kimbrough et al., 2017; Lindell et al., 2017; Loi et al., 2010; Melendez, 2011; Rosenwasser et al., 2013; Simms et al., 2008, 2010; Spoelder et al., 2015). Additionally, 2BC home cage drinking causes a gradual increase in ethanol intake over the course of weeks, often thought to reflect a clinically relevant binge drinking phenotype, which is augmented in IA2BC compared to a continuous 2BC procedure (Lindell et al., 2017; Loi et al., 2010; Rosenwasser et al., 2013; Simms et al., 2008, 2010; Spoelder et al., 2015). Importantly, these features increase the translational relevance of IA2BC by reflecting cycles of binge and withdrawal observed in the clinical population (Bowen et al., 2022; Carnicella et al., 2014; Spear, 2020).

While evidence suggests that IA2BC has superior face validity to continuous 2BC paradigms, subjects in both paradigms typically retain *ad libitum* access to ethanol. Thus, while the access schedule differs *between* drinking sessions, both paradigms have continuous access (ContA) to ethanol *within* a session. While useful, this does not reflect the dynamic settings and associated factors that can drive variable rates of within-session consumption, and related craving, in the clinical population (Dvorak et al., 2014; Fischer et al., 2023; Fridberg et al., 2021; King et al., 2025; Pelloux et al., 2019; Prignitz et al., 2024; Wray et al., 2014). Clinical research makes it increasingly clear that the rate of ethanol consumption within a drinking session, not just the total amount, has clinical utility for informing risk of AUD development and severity of AUD symptoms (Carpenter et al., 2017, 2019; Gowin et al., 2017; Simons et al., 2015; Sloan et al., 2020). As such, within-session intermittent access (IntA), defined as providing discrete reinforcer access periods interspersed with periods of no reinforcer access within a given drinking session, may serve to better model dynamic settings and drinking patterns associated with AUD that are observed in clinical populations (Ardinger et al., 2022; Leeman et al., 2010).

While not studied extensively in the context of ethanol consumption, within-session IntA for cocaine and opioids consistently elicits greater reward seeking, increases cue-induced reinstatement, and heightens SUD-related phenotypes including escalation and aversion-resistant intake compared to ContA schedules (often denoted as long access [LgA]) (Allain & Samaha, 2019; Beasley et al., 2023; Calipari et al., 2013, 2015; Calipari & Jones, 2014; D’Ottavio et al., 2023; Fragale et al., 2020; James et al., 2019; Nicolas et al., 2019; Rakowski et al., 2025; Samson et al., 2022; Singer et al., 2018). This effect is thought to be driven, in large part, by dynamic, fluctuating drug concentrations and full clearance of drug between intoxication sessions, which mimic human patterns of binge intake (Allain & Samaha, 2019; Kawa et al., 2019; Zimmer et al., 2011, 2012). These data suggest that within-session IntA may also be able to promote increased ethanol consumption and exacerbate AUD-relevant phenotypes. However, this possibility has not been widely assessed due to the fact that, unlike psychostimulants and opioids, ethanol is metabolized at much slower rates (Wilson & Matschinsky, 2020), precluding the ability of within-session IntA to drive dynamic, fluctuating blood ethanol concentrations (BECs) thought to be required for these effects. However, recent work found that within-session IntA for non-drug reinforcers engenders more robust reward seeking, cue-induced reinstatement, and higher motivation than within-session ContA procedures (Beasley et al., 2022; Muñoz-Escobar et al., 2019; M. J. F. Robinson et al., 2023). Interestingly, this effect was greater when periods of reinforcer access and interval were unpredictable (M. J. F. Robinson et al., 2023). These data imply that the effects of within-session IntA drug cannot be fully explained by reinforcer pharmacokinetics, suggesting that these effects may in fact generalize to ethanol. In line with this, two studies have shown that within-session IntA to ethanol promotes higher voluntary intake, ethanol seeking, and aversion-resistant seeking compared to within-session ContA (Gage et al., 2023; Tomie et al., 2006).

Altogether, these data imply that within-session IntA, and unpredictable IntA (uIntA) in particular, to ethanol could exacerbate ethanol intake and AUD-related symptomatology to a greater degree than within-session ContA paradigms. However, prior work has yet to examine the effect of uIntA on ethanol consumption or AUD-related phenotypes associated with ethanol dependence, including escalation of ethanol consumption and aversion-resistant drinking. As such, the present study sought to examine voluntary ethanol consumption prior to and following the induction of ethanol dependence under an uIntA schedule and directly compare this to a ContA schedule. We hypothesized that uIntA would promote higher ethanol intake and patterns of consumption that drove higher within-session BECs prior to the development of dependence than ContA. Furthermore, we hypothesized that uIntA would exacerbate dependence-induced escalation and aversion-resistant intake compared to ethanol-dependent rats drinking under a ContA schedule.

## Methods

### Animals

Adult male and female Long-Evans rats (Envigo, Indianapolis, IN) were utilized for all experiments (n=24/sex). Rats were ∼P60 on arrival and allowed one week of acclimation in the vivarium prior to testing. Rats were singly housed in standard Plexiglas cages with corncob bedding (‘Bed o’ Cobs’, The Andersons Inc.) in a temperature-controlled room under a 12:12 hr reverse light/dark cycle (lights on at 22:00). Rats were provided *ad libitum* access to standard enrichment, chow (Teklad 7912, Envigo), and water throughout the duration of the study. All experimental procedures were approved by the University of Illinois Chicago Institutional Animal Care and Use Committee and adhered to the NIH Guidelines for the Care and Use of Laboratory Animals.

### Schedules of ethanol access

Rats were trained to drink ethanol (20% v/v) in 1 hour drinking sessions using a lickometer-equipped apparatus (11.63”X9.78”X7.35”) with a rod floor and custom-made 3D-printed pans for waste collection, which was housed in a sound-attenuating cubicle (25”X17.5”X23.5”; MED Associates). For all sessions, background noise and ventilation within the apparatus was provided by a fan that remained on throughout testing. Ethanol was provided using a retractable sipper consisting of a Lixit spout and 8 fl. oz. bottle (MED Associates, ENV-252M). Drinking patterns were assessed using time-stamped lickometry and total ethanol intake was measured via change in bottle weight before and after the session. Bottle testing was conducted to assess spillage as a result of bottle placement, testing, and removal. An average spillage of 0.025 g change in bottle weight (range: 0-0.06 g) was identified, equating to ∼4 mg of ethanol lost (range:0-9.88 mg). Importantly, spillage was not significantly different when bottles were tested using ContA or uIntA (ContA:4.174±0.8488, uIntA:3.954±0.6783). To control for the effect of this spillage, we deducted the maximum amount of spillage identified (0.06 g change in bottle weight equating to 9.88 mg of ethanol) from all of our intake calculations prior to analysis. Upon completion of each session, the area underneath the bottle was inspected for fluid leakage. Sessions where significant leakage was identified (n=3 of 1,680) were excluded from all analyses. All testing was performed during the dark (active) phase of the light-dark cycle and rats were protected from ambient lighting during transfer to the testing room using custom-made black-out covers. Once in the experimental room, red light was used to remove, weigh, and transfer rats into the testing apparatus. Between sessions, boxes and rod floors were thoroughly cleaned using 70% ethanol, allowed to dry, and fitted with a clean waste pan. Rod floors and waste pans were disinfected daily using *Conflikt* after all testing was completed.

For the first drinking session, all rats experienced a continuous access (ContA) schedule. After the first session, rats were randomly assigned to either ContA or unpredictable intermittent access (uIntA) schedules and experienced the same schedule for the remainder of testing. ContA consisted of 60 minutes of uninterrupted access to the ethanol-containing sipper (**Fig. 1A**). uIntA consisted of sipper access spread across 26 epochs of variable duration (15s, 20s, 30s, 40s, 60s, 90s, 120s) and interval (30s, 60s, 2.5 min, 5 min, 10 min) (**Fig. 1B**). Sipper access in both schedules was accompanied by house light illumination (producing ∼20-25 lux at the center of the apparatus), which served as a discriminative stimulus (DS) signaling ethanol availability. For uIntA sessions, DS and interval (or inter-trial interval [ITI]) durations occurred a fixed number of times per session, as described in **Table 1**. Individual DS and ITI durations were pulled randomly without replacement from the list of all possible durations within each session, leading to a pseudorandom order of DS presentations (or ethanol access periods) that amount to 15 min per session.

**Figure 1.**
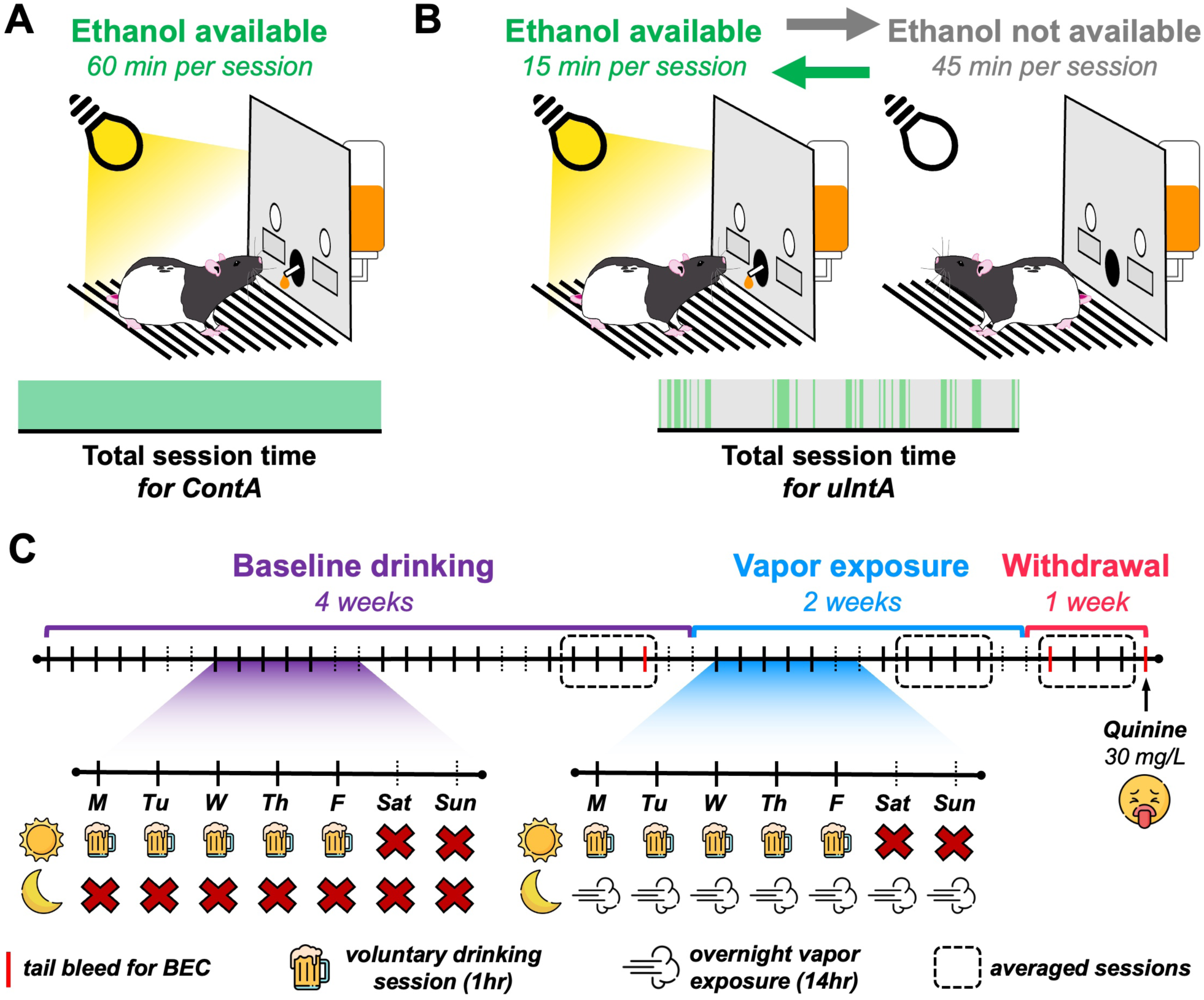
Experimental design. **(A)** Rats that underwent daily continuous access (ContA) sessions were allowed 60 min of uninterrupted ethanol access paired with house light illumination. **(B)** Rats that underwent daily unpredictable intermittent access (ulntA) sessions experienced variable periods of ethanol availability (amounting to 15 min) signaled by house light illumination. separated by variable periods without ethanol access (amounting to 45 min). **(C)** Rats underwent a total of seven weeks of voluntary drinking sessions. After a four-week baseline period, rats underwent two weeks of chronic intermittent ethanol or air vapor exposure during which drinking sessions continued Mon-Fri. Drinking sessions continued into withdrawal after completion of vapor exposure. On the final day of withdrawal testing, ethanol was adulterated with 30 mg/L quinine to assess aversion-resistant drinking. For each experimental timepoint, data was averaged across the final four sessions (indicated by dotted squares)

**Table 1.** Discriminative stimulus (ethanol access) and intertrial interval (no ethanol access) periods within the uIntA schedule.

|  | Duration<br>(s) | # of<br>occurrences | Total time<br>(min) |
| --- | --- | --- | --- |
| **Discriminative**<br>**Stimulus (DS)**<br>*(ethanol access)* | 15 | 10 | 2.5 |
|  | 20 | 5 | 1.67 |
|  | 30 | 2 | 1 |
|  | 40 | 2 | 1.33 |
|  | 60 | 5 | 5 |
|  | 90 | 1 | 1.5 |
|  | 120 | 1 | 2 |
| ..... |  |  |  |
| **Intertrial**<br>**Interval (ITI)**<br>*(no ethanol access)* | 30 | 11 | 5.5 |
|  | 60 | 7 | 7 |
|  | 150 | 3 | 7.5 |
|  | 300 | 3 | 15 |
|  | 600 | 1 | 10 |

### Chronic intermittent ethanol vapor exposure

After four weeks of daily ethanol drinking sessions, rats were rendered dependent using a standard 14-day chronic intermittent ethanol vapor exposure procedure as previously described (Glover et al., 2019, 2021; Przybysz et al., 2024; Ramirez et al., 2024). Rats experienced either vaporized ethanol (EtOH-exposed group) or room air (Air-exposed group) for 14 hr/day (18:00-8:00) for 14 consecutive days. Behavioral signs of intoxication were assessed daily in EtOH-exposed rats using a previously published subjective rating scale (Glover et al., 2021; Ramirez et al., 2024) ranging from 1 (no signs of intoxication) to a 5 (total loss of consciousness). Blood samples (40 uL) from EtOH-exposed rats were extracted immediately upon removal from chambers via tail vein nick on days 2, 6, 10, and 14 (±1 day) of the exposure period for the measurement of blood ethanol concentration (BEC) as a result of ethanol vapor exposure. Air-exposed counterparts were administered a tail pinch on the same schedule to mimic the temporary discomfort evoked by blood collection experienced by intoxicated rats.

### Timeline of experiments

A schematic describing the experimental timeline is shown in **Fig. 1C**. Voluntary ethanol drinking sessions occurred daily, excluding weekends, throughout all experimental timepoints for a total of seven weeks. Baseline (BL) data was collected during the first four weeks of testing. Daily drinking sessions continued during the day while rats underwent vapor exposure overnight. This was followed by one week of data collection during withdrawal (WD) after vapor exposure was completed. On the final day of testing (WD day 5), ethanol was adulterated with quinine (30 mg/L) to assess aversion-resistant drinking. Blood samples (40 uL) were collected from all rats immediately following completion of the last baseline, first WD, and last WD drinking sessions to measure [actual] blood ethanol concentration (aBEC) achieved as a result of voluntary ethanol consumption.

### Blood ethanol concentration

Upon collection, blood samples were immediately centrifuged (10,000 rcf, 10 min, 4°C). Plasma supernatant (20 uL) was aliquoted into 0.5 mL microcentrifuge tubes and stored at −20°C. BECs was measured using an Analox Alcohol Analyzer (Analox Instruments Ltd, United Kingdom) within 30 days of sample collection.

Estimated blood ethanol concentration (eBEC) was calculated at each experimental timepoint using a modified application of the commonly described Widmark equation paired with a zero order model of ethanol clearance (Widmark, 1981):

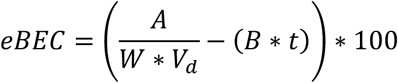

where *A* is defined as the amount of ethanol (g) consumed, *W* indicates the weight (kg) of the rat, *t* is defined as the amount of time (hr) since the start of drinking, *V_d_* is a constant (L/kg) that informs the volume diffusion of ethanol in the body (denoted as *R* in the standard Widmark application), and *B* is defined as the barrier term that simulates linear removal of available ethanol (g/L/hr; denoted as a β in the standard Widmark). In the standard use of the Widmark equation, the linear removal term is thought to exclusively reflect the linear portion of hepatic metabolism occurring following complete absorption of ethanol into the bloodstream (Jones, 2019; Posey & Mozayani, 2007). This is an especially important and critical assumption in medicolegal contexts where the only ethanol consumption information likely known is the total number of drinks consumed and an approximate time when drinking ended. In our modified use of this equation, the barrier term (*B*) consistently ‘removes’ a defined amount of ethanol consumed by the animal at every timepoint calculated to attempt to model any process that serves as a barrier to ethanol availability in the blood of animals that are freely consuming. This includes, but is not necessarily limited to, first pass metabolism in the gastric compartment, gastric emptying rate and absorption into the portal vein, first pass metabolism in the hepatic compartment, and the standard linear and non-linear portions of ethanol metabolism in the hepatic compartment (Goldman et al., 2025; Jones, 2019; Wilson & Matschinsky, 2020). The final eBEC output from this equation was multiplied by 100 to convert from g/L to mg/dL (i.e. mg%) for all calculations.

For these experiments, *V_d_* and *B* were individually defined for each rat to best map onto ‘ground truth’ end-of-session aBEC values obtained across the experimental timeline (**Fig 1C, Supp. Fig 1A-B**). Individual estimation of these values allows us to account for individual variability in factors affecting the volume distribution of ethanol as well as the gastric emptying, first pass gastric and hepatic metabolism, and systemic metabolism of ethanol in our free-feeding, outbred Long-Evans rats over the experimental timeline (Goldman et al., 2025; Jones, 2019; Wilson & Matschinsky, 2020). To ensure biological relevance of these measures, individual estimation of *V_d_* was constrained to sex-defined ranges of volume diffusion of ethanol provided by published literature in male and female Long-Evans rats (B. Webb et al., 2002). Individual estimation of *B* was constrained in a similar fashion by values describing the linear portion of ethanol metabolism (B. Webb et al., 2002) but was allowed to vary outside of this range in certain instances (described below) since, in our application, *B* is not likely to reflect linear ethanol metabolism exclusively. To determine individually defined *V_d_* and *B* values, end-of-session eBECs were simulated across a range of values corresponding to *V_d_* and *B* for each session where aBECs were measured. For *V_d_*, values tested ranged from 0.5-0.9 L/kg in 0.01 increments. For *B*, values tested ranged from 0.1-1.0 g/L/hr in 0.01 increments. For each session where tail blood was collected, end-of-session eBEC values produced from the combination of *V_d_* and B values across the stated ranges were compared to the end-of-session aBEC value obtained from tail blood. For each session, sets of *V_d_* and B pairs that produced a residual between end-of-session eBEC and aBEC values of less than 20 mg/dL and fell within ± 2 standard deviations (SD) of sex-defined ranges of these values in the literature (B. Webb et al., 2002) were averaged to generate a ‘best’ *V_d_* and *B* pair. If no *V_d_* and *B* pairs were identified where both these criteria were met, the *V_d_* value was retained within the sex-defined range while *B* was adjusted to produce a residual of less than 20 mg/dL. All *V_d_* and *B* pairs meeting these criteria were averaged in order to delineate a ‘best’ *V_d_* and *B* pair for that session. As such, the *B* values for some rats varies beyond ± 2 SD from the range of values describing the linear portion of ethanol metabolism provided by the literature (B. Webb et al., 2002). Because aBECs of ≤20 mg/dL are considered within the range of error using the Analox Analyzer, any sessions where intake was ≤0.1 g/kg and the corresponding aBEC was ≤20 mg/dL were excluded from these estimates. After these considerations, the resultant ‘best’ *V_d_* and *B* pairs established for each session where tail blood was collected were then averaged to obtain a global individually defined value for each animal, which was then utilized to calculate individualized eBECs for all timepoints. Individual estimation of these values helps better control for variability observed in these processes across animals as well as controlling for possible changes in these processes across our experimental timeline, including the possible development of metabolic tolerance after ethanol vapor exposure (Cederbaum, 2012; Glover et al., 2021; Goldman et al., 2025; Jones, 2019), **Supp. Fig 1B**). For *A,* the total number of licks was divided by the total ethanol intake (in g) for each session to obtain a “gperlick” measurement that was used to calculate the total amount of ethanol consumed based on the number of licks performed at any given timepoint within the session. eBEC curves throughout the session were then calculated from the accumulation of ethanol intake via this “gperlick” measurement in 1 min bins. Iterative calculation of eBECs using this method is visualized in **Supp. Fig 1C**. It is important to note that the range of possible eBECs achievable for a given animal in a given session is governed by their individually estimated *V_d_* and *B* values, as well as the total intake observed. The final within-session eBEC estimates in this range are then governed by the pattern of drinking observed via lickometry (**Supp. Fig 1C**).

### Lick microstructure analysis

Lick microstructure analysis was utilized as described previously (Hou et al., 2024; Patwell et al., 2021; Pitock et al., 2025; Wheeler et al., 2025). In brief, bouts within a session were defined as three or more licks with an interlick interval of less than 1 second. Importantly, periods when ethanol was not available were subtracted from uIntA analyses, allowing us to analyze lick microstructure in a pseudo-consecutive 15-minute (900 sec) period of ethanol availability. Bout probability for each analyzed session was defined by delineating bout “on” (1)/ bout “off” (0) periods during ethanol availability at the sampling rate of our lickometer-equipped apparatus (100 Hz; 1 sample/10 ms). Bout probability timeseries were then averaged across the final four sessions within each timepoint for each animal. To enable direct comparisons between ContA and uIntA schedules, bout probability was binned into 30 s periods of ethanol availability, leading to asymmetrical numbers of datapoints between schedules (ContA [120 datapoints amounting to 3600 s]; uIntA [30 datapoints amounting to 900 s]). Bout probability analyses that were limited to DS presentation periods in uIntA rats only consisted of data binned in 1 s periods of DS presentation time, leading to asymmetrical datapoints for every DS duration.

### Statistical analysis

Unless otherwise indicated, data from the final four voluntary drinking sessions at each timepoint were averaged together for each rat for statistical analyses in order to ensure that data from each time point was equally represented in group analyses. Data were pooled across sexes for all analyses due to insufficient power to determine significant effects of sex in the current experiments. Rats exhibiting an average intake of <0.1 g/kg at baseline (n = 2-3/group) were excluded due to insufficient ethanol consumption. Importantly, there was no significant difference in the proportion of rats excluded in either schedule of access (Fisher’s exact test, *p*>0.9999). Upon collection of all experimental data, statistical outlier analyses (ROUT) were performed on all datasets with a False Discovery Rate (Q) of 0.02%.

For all analyses related to bout probability, linear mixed effects models (LMEs) were utilized due to the asymmetrical nature of the data structure. For symmetrical data structures, multifactorial ANOVAs or repeated measures (RM) ANOVAs were utilized where appropriate. Comparisons of group differences after baseline were performed using omnibus multifactorial RM ANOVAs or LMEs with time, schedule, and vapor exposure as primary independent variables. This was followed by two-way RM ANOVAs or LMEs to further identify effects of vapor and schedule on dependence-induced escalation of ethanol consumption and aversion-resistant drinking. LMEs were also utilized to examine the effect of multiple parameters on current DS bout probability during DS presentations in the uIntA group including: (1) time within the current DS, (2) latency of DS onset from session start, (3) duration of prior ITI, (4) experience of a termination of bout by DS offset during the prior DS, (5) amount of EtOH consumed (in g/kg) during the prior DS, and (6) cumulative EtOH consumed prior to the current DS. For baseline sessions, a single model incorporating all of the above parameters was utilized. For sessions occurring after baseline, data from the final four sessions of vapor exposure and withdrawal (8 sessions total) were pooled to assess the interaction of vapor exposure with all of the above parameters. Due to the nature of these parameters, data from the first DS presentation were removed for analysis. For parameters related to ethanol consumption, the amount of ethanol consumed was z-scored across all individual DS presentations for analysis. For data structures where RM ANOVAs and LMEs were not used, unpaired t-tests/Mann-Whitney tests were utilized as appropriate. Spearman’s correlation was utilized for all correlational analyses. A ≥20% change criterion was utilized to classify animals as escalators and quinine sensitive. This criterion allowed for accurate identification of animals exhibiting a change in drinking phenotype while accounting for minor day-to-day fluctuations in intake commonly observed in individual animals. Fisher’s exact test was utilized to determine significant differences in the proportion of rats exhibiting specific phenotypes.

All analyses, except for linear mixed effect modeling, were conducted using Prism 10 (Graphpad). LMEs were generated via the *fitlme()* function in MATLAB (R2024a, MathWorks). Composite hypothesis testing was conducted utilizing the *anova()* function. The Satterthwaite method for designation of degrees of freedom was utilized due to relatively large sample sizes. The significance threshold for all experimental tests was set at *p* < 0.05. Data in **Supp Table 1** presents test statistics derived from traditional methods of analysis and data in **Supp Table 2** provides test statistics derived from LMEs, including beta estimates and standard error, that are not reported in-text. Error bars for all graphs and in-text reports reflect standard error of the mean (SEM). For all graphs, ContA is depicted with solid lines and closed datapoints, whereas uIntA is depicted with dotted lines and open datapoints, unless otherwise described. Circles indicate data from male rats and triangles indicate data from female rats.

## Results

### Validation of estimated blood ethanol concentration (eBEC) analysis

Prior work has shown that the rate of ethanol consumption, and resultant pharmacokinetics, can predict risk for binge drinking and development of AUD (Carpenter et al., 2017, 2019; Gowin et al., 2017; Simons et al., 2015; Sloan et al., 2020). Clinical assessments often use eBECs to identify how ethanol pharmacokinetics relate to the subjective effects of ethanol, relative intoxication, and expression of AUD symptomatology (Hultgren et al., 2025; Piasecki, 2019). Surprisingly, eBEC analyses have not been widely adopted in preclinical assessments despite its translational value and ability to provide a deeper understanding of how ethanol intake patterns, and resultant eBEC dynamics, drive AUD-relevant symptomatology (Grahame, 2026). As such, we sought to develop an eBEC analysis pipeline to allow for within-session estimation of BECs in animals voluntarily consuming ethanol at variable rates across our experimental timepoints.

BECs measured from blood samples obtained at the termination of three separate drinking sessions across the experimental timeline (**Fig. 1C**), individually defined estimates of the volume of distribution for ethanol (*V_d_*), individually estimated barrier terms (*B*), and ‘mgperlick’ values were used in combination to calculate eBECs using a modified application of the Widmark equation (**Fig 2A-B**). Individually defined *V_d_* and *B* estimates were determined as the values that produced the smallest residual between final eBEC and aBEC of the drinking sessions where tail blood was collected (**Supp. Fig 1A-B**). Importantly, *V_d_* values were retained within the physiological range reported in the prior literature (Aull et al., 1956; Chung et al., 2008; Crippens et al., 1999; Doherty & Gonzales, 2015; Kharbouche et al., 2010; Leasure & Nixon, 2010; Little et al., 1996; Varlinskaya & Spear, 2004; B. Webb et al., 2002). While *B* varied outside of the range of values reported in the literature for the linear portion of hepatic metabolism (Aull et al., 1956; Chung et al., 2008; Crippens et al., 1999; Doherty & Gonzales, 2015; Kharbouche et al., 2010; Leasure & Nixon, 2010; Little et al., 1996; Varlinskaya & Spear, 2004; B. Webb et al., 2002), this is expected since *B* is likely to reflect all pharmacokinetic processes that may serve as a barrier to ethanol availability in the blood, including linear hepatic metabolism. Inter-lick interval distributions for all sessions where eBECs were calculated peaked at ∼140 ms and were not different between schedules (**Fig 2C**), supporting the fidelity of lick timestamps in reflecting biologically relevant drinking in agreement with prior work (Lin et al., 2013). A significant positive correlation was observed between total number of licks and total intake within each session (**Fig 2D**), confirming the fidelity of the lickometer-equipped apparatus to accurately reflect intake throughout the session. Importantly, no significant difference was identified between schedules in the ‘mgperlick’ measurement used to calculate ethanol intake across the session (**Fig 2E**). Furthermore, there was no significant effect of schedule on individual *V_d_* estimates (**Fig 2F**), *B* estimates (**Fig 2G**), or residuals (**Fig 2H**) identified using these values. In addition, no significant between group differences were observed in *V_d_* estimates, *B* estimates, average residual, or calculated mg of EtOH per lick when data were examined by ethanol vapor group in addition to schedule (**Supp Table 1**). Spearman’s correlation revealed a significant positive correlation between intake and ‘end of session’ aBEC measured for sessions where tail blood was collected (**Fig 2I**). We also observed significant positive correlations between ‘end of session’ eBECs and corresponding aBECs across sessions where tail blood was collected (**Fig 2J**) as well as between peak eBEC and intake achieved at each experimental timepoint (**Fig 2K**). These data confirm that calculations used to estimate within-session eBECs accurately reflect aBECs achieved in each rat at the end of the session and are scaled appropriately with the within-session intake achieved by each rat. Altogether, these data support the validity of the current eBEC calculation to accurately reflect aBECs achieved as a result of voluntary ethanol drinking throughout a session using a lickometer-equipped apparatus.

**Figure 2:**
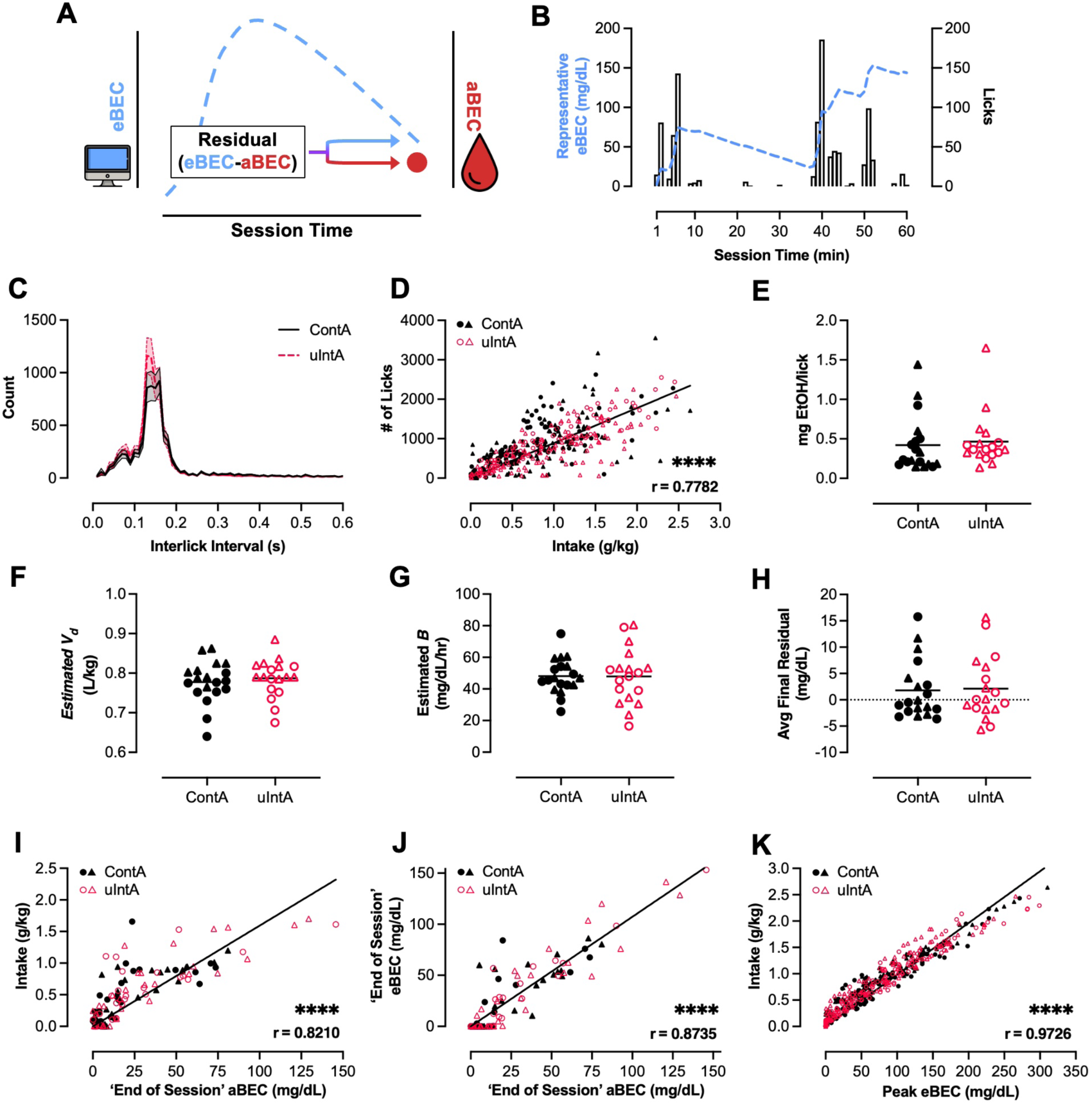
Validation of estimated blood ethanol concentration (eBEC) analysis. **(A)** eBECs were calculated and validated with end-of-session aBECs. **(B)** Representative eBEC curve (blue line) plotted with number of licks occurring within each minute of the session (black bars) to exemplify use of ‘mgperlick’ measurement in eBEC calculation. **(C)** Inter-lick interval distributions for both ContA and ulntA schedules. **(D)** Correlation between total number of licks and total intake for all sessions. Amount of EtOH consumed (in mg) per lick **(E),** individually defined estimated Widmark constants *(R;* L/kg) **(F),** individually defined estimated β ethanol elimination rates **(G),** and average residual [‘end of session’ eBEC - ‘end of session’ aBEC] **(H)** for ContA and ulntA rats. (I) Correlation between average intake and average ‘end of session’ aBEC for drinking sessions where tail blood was collected. **(J)** Correlation between ‘end of session’ eBEC and ‘end of session’ aBEC for drinking sessions where tail blood was collected. **(K)** Correlation between peak eBEC and intake across experimental timepoints.

### Intoxication as a result of chronic intermittent ethanol vapor exposure

Average intoxication scores during chronic intermittent ethanol vapor exposure were not significantly different between schedules (ContA: 2.086±0.059, uIntA: 2.087±0.069, t(17)=0.01755, *p*=0.9862). In addition, there was no significant effect of drinking schedule on BECs achieved during ethanol vapor exposure (ContA: 259.9±16.63 mg/dL, uIntA: 254±21.71 mg/dL, t(17)=0.2206, *p*=0.8280). As expected, a significant positive correlation was observed between intoxication score and BECs achieved as a result of vapor exposure (r=0.6914, *p*<0.0001). Altogether, these data corroborate prior work (Glover et al., 2019, 2021; Przybysz et al., 2024; Ramirez et al., 2024) and demonstrate a similar degree of intoxication during chronic intermittent ethanol vapor exposure between rats experiencing ContA or uIntA.

### Schedule of access does not affect drinking dynamics prior to ethanol dependence

To determine whether uIntA can increase ethanol consumption, and other relevant parameters, compared to ContA, we compared overall intake between groups prior to ethanol dependence. As shown in **Fig. 3**, rats that underwent uIntA drinking exhibited similar levels of ethanol consumption at baseline as rats that underwent ContA drinking. Indeed, analysis of intake across all pre-vapor sessions as well as average intake during the final four baseline sessions found no significant main effect of schedule on intake (**Fig. 3A**). Moreover, aBECs captured after the final baseline drinking session were not significantly different between schedules (**Supp Table 1**). In agreement with this, uIntA and ContA groups also exhibited similar within-session eBEC dynamics (**Fig 3B**) and bout probability (**Fig 3C, Supp Fig 2A-B**) at baseline. Importantly, differences in these measures remained nonsignificant when analyzed by future vapor group assignment (**Fig 3D-F**). Furthermore, Fischer’s exact test identified that the proportion of rats whose average baseline intake was classified as light (<0.5 g/kg), moderate (0.5-1.0 g/kg), or heavy (>1 g/kg) did not differ between schedules or vapor exposure groups (**Fig 3G**-**H**). Altogether, these data show that uIntA engenders moderate levels of voluntary ethanol consumption to a similar degree as ContA.

**Figure 3.**
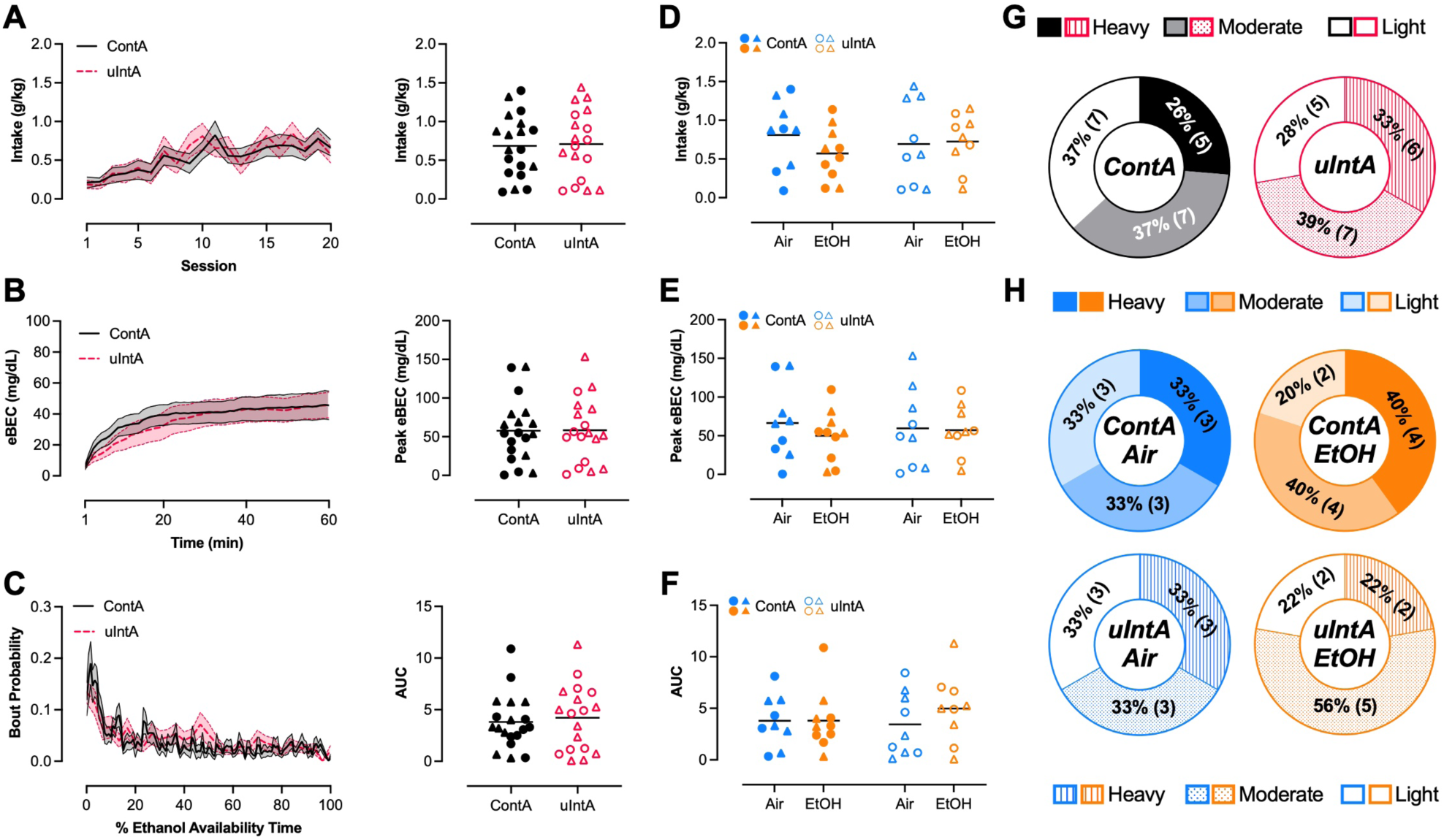
Schedule of access does not affect ethanol consumption, eBEC dynamics, or drinking patterns at baseline. **(A)** Daily ethanol intake across all baseline drinking sessions (left) and average intake from the final four sessions (right) of baseline prior to vapor exposure. **(B)** Average within-session eBECs (left) and average peak eBEC (right) at baseline. **(C)** Average within-session bout probability at baseline (left). Ethanol availability time was normalized to 60 min (from 15 min) for the ulntA group to enable direct comparison with the ContA group. Area under the curve (AUC) of baseline within-session bout probability (right). Average ethanol intake **(D),** peak eBEC achieved **(E),** and AUC of bout probability **(F)** at baseline separated by schedule and vapor group assignment. The proportion of rats classified as light (< 0.5 g/kg), moderate (0.5 g/kg - 1 g/kg), or heavy (> 1 g/kg) drinkers at baseline grouped by schedule only **(G)** and by schedule and vapor group assignment **(H).** Error bars indicate ± SEM. Solid lines and closed datapoints indicate ContA. Dashed lines and open datapoints indicate ulntA. Circles and triangles indicate males and females, respectively. n=4-6/sex/group.

### Dependence-induced escalation of ethanol consumption differs across schedules

Although we expected that all rats exposed to chronic intermittent ethanol vapor would exhibit escalated ethanol intake, we hypothesized that uIntA would exacerbate escalation relative to ContA counterparts. To assess this, we performed omnibus multifactorial RM ANOVAs or LMEs utilizing time (continuous or categorical [i.e. timepoint]), schedule (ContA vs. uIntA), and vapor (Air vs. EtOH) as independent variables across all relevant dependent variables. As denoted in **Supp Table 1 & 2**, several analyses indicate trending three-way interactions and significant two way-interactions of schedule and vapor, suggesting non-independent effects of both factors on each dependent measure. As such, we conducted RM two-way ANOVAs or LMEs to determine the effect of vapor within each schedule (ContA Air- vs. EtOH-exposed; uIntA Air- vs. EtOH-exposed). Separate analyses were performed to determine the effect of schedule within ethanol dependent rats (ContA EtOH-exposed vs. uIntA EtOH-exposed).

### Dependence-induced escalation is transient in ContA drinking rats

When drinking is examined across sessions, no significant main effect of vapor or vapor by timepoint interaction was identified in rats that drank ethanol under a ContA schedule (**Fig. 4A**). However, a significant vapor by timepoint interaction was detected when intake was assessed during the final four sessions of each timepoint (**Fig. 4B**). Tukey’s posthoc comparisons indicate that EtOH-exposed ContA rats drank significantly more ethanol during vapor exposure compared to baseline (*p*=0.0194) and withdrawal (*p*=0.0347). Additionally, EtOH-exposed rats drank more during vapor exposure than Air-exposed counterparts (*p*=0.0229). This difference was also apparent when data were assessed in terms of change in intake relative to baseline (**Fig. 4C**). Šidák’s posthoc comparisons found that EtOH-exposed ContA rats increased their ethanol intake to a greater degree during vapor exposure than did Air-exposed ContA rats (*p*=0.0140). Furthermore, EtOH-exposed rats exhibited a greater increase in intake during vapor exposure than during withdrawal (*p*=0.0012). In corroboration with average intake increasing during vapor exposure, we also observed a significant main effect of vapor group and vapor by time interaction on eBECs achieved during daily drinking sessions occurring throughout the 14d vapor exposure paradigm (**Fig. 4D**). However, Šidák’s posthoc comparisons did not reveal any significant simple main effects. Moreover, this effect of vapor group did not persist into withdrawal (**Fig. 4E**). This is corroborated by a significant vapor by timepoint interaction on average peak eBEC achieved during the final four sessions at each timepoint (**Fig. 4F**). Tukey’s posthoc comparisons indicate the EtOH-exposed ContA rats achieved higher peak eBECs during vapor exposure compared to baseline (*p*=0.0105) and withdrawal (*p*=0.0210). Moreover, EtOH-exposed ContA rats achieved a higher average peak eBEC than Air-exposed counterparts during drinking sessions occurring over the course of vapor exposure (*p*=0.0049). In agreement with eBECs, aBECs were not significantly different between Air- and EtOH-exposed rats at 24 hr withdrawal (t(17)=2.026, *p*=0.0587; Air: 25.63±8.57 mg/dL; EtOH: 8.640±2.02 mg/dL). This transient increase in intake and intoxication is further corroborated by changes in bout probability, such that a main effect of vapor and a vapor by time interaction was detected during drinking sessions occurring during vapor exposure (**Fig. 4G, Supp Fig 2C,E**). However, these effects are not retained during withdrawal (**Fig. 4H, Supp Fig 2D-E**). Analysis of bout probability area under the curve (AUC) found a significant vapor by timepoint interaction with Tukey’s posthoc comparisons revealing that EtOH-exposed ContA rats had significantly greater AUC during the vapor timepoint than the withdrawal timepoint (**Fig. 4I**; *p*=0.0449). While not significant, EtOH-exposed ContA rats exhibited a trend for greater AUC during the vapor exposure timepoint compared to baseline (*p*=0.0785). Additionally, EtOH-exposed ContA rats exhibited significantly greater AUC than Air-exposed counterparts during the vapor exposure timepoint (*p*=0.0091). Finally, Fischer’s exact test identified that the proportion of rats defined as escalators (exhibiting ≥20% increase in intake from baseline) was significantly higher in EtOH- than Air-exposed rats during drinking sessions that occurred in parallel with vapor exposure (**Fig. 4J**; *p*=0.0027) but not during withdrawal (**Fig. 4K**; *p*=0.3133). Altogether, these data indicate that ethanol-dependent rats under a ContA schedule exhibit a transient escalation of ethanol intake observed only during chronic intermittent ethanol vapor exposure.

**Figure 4.**
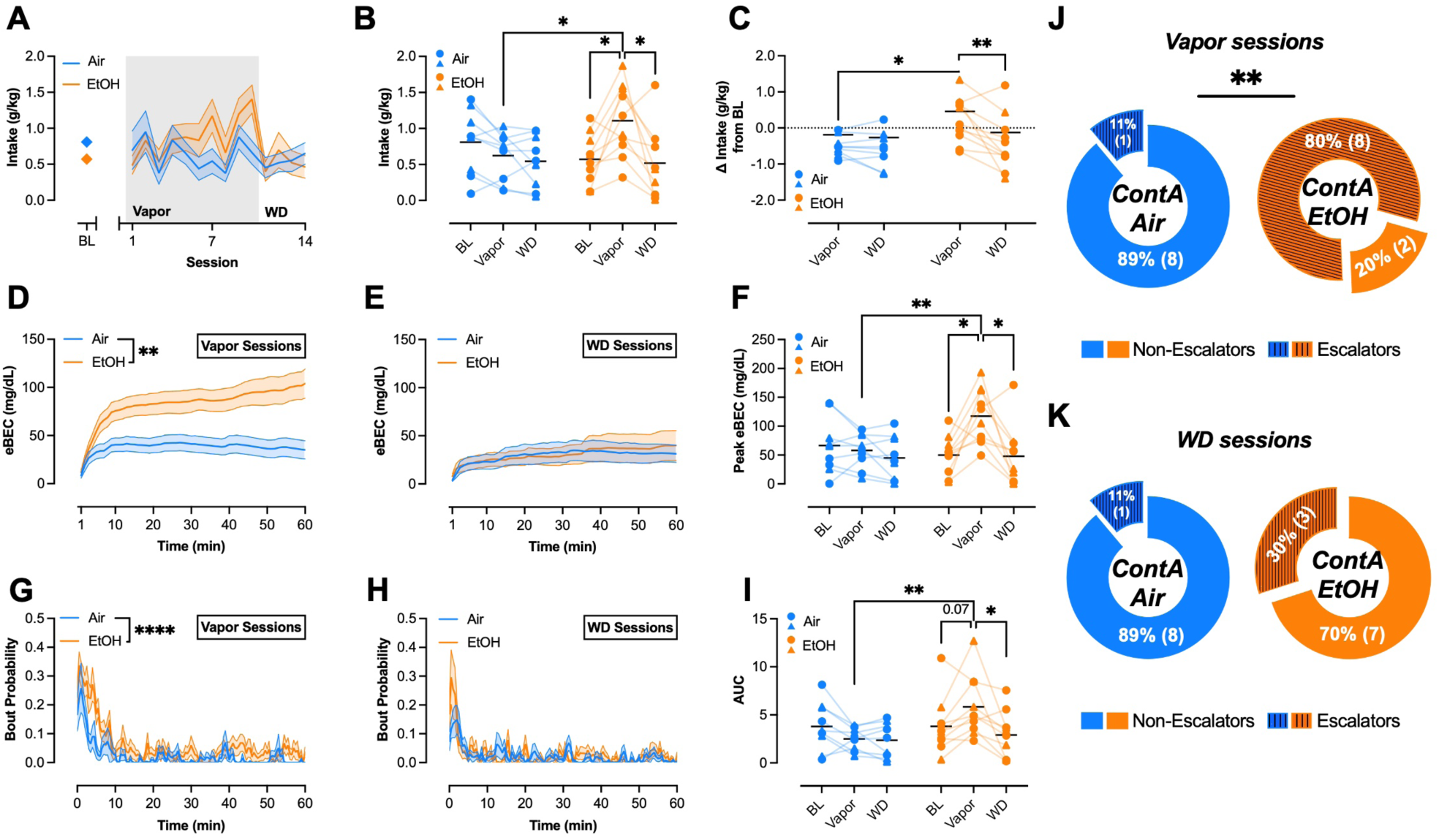
Dependence promotes transient escalation of ethanol consumption in rats drinking under a ContA schedule. **(A)** Daily ethanol intake during vapor exposure and withdrawal timepoints relative to average intake during the final four baseline sessions. **(B)** Average intake across all timepoints. **(C)** Change in intake from baseline during vapor exposure and withdrawal. **(D)** Average within-session eBECs achieved in drinking sessions that occurred during vapor exposure. **(E)** Average within-session eBECs achieved in drinking sessions during withdrawal. **(F)** Average peak eBEC achieved across all timepoints. **(G)** Average within-session bout probability during vapor exposure. **(H)** Average within-session bout probability during withdrawal. (I) Area under the curve of bout probability across all timepoints. Proportion of Air- and EtOH-exposed ContA rats defined as escalators (≥ 20% increase from baseline) or non-escalators during vapor exposure **(J)** and withdrawal sessions **(K).** Error bars indicate ± SEM. Solid lines and closed datapoints indicate ContA. Circles and triangles indicate males and females, respectively. n=4-6/sex/group.

### Dependence-induced escalation is sustained in uIntA drinking rats

In rats that underwent uIntA drinking, a significant main effect of vapor was detected in per session intake (**Fig. 5A**), indicating that EtOH-exposed uIntA rats drank more during all post-baseline sessions compared to Air-exposed counterparts. This is reinforced by a significant vapor by timepoint interaction observed in average intake during the final four sessions of each timepoint (**Fig. 5B**). Tukey’s posthoc comparisons indicate that EtOH-exposed uIntA rats exhibited significantly higher intake during the vapor (*p*=0.0168) and withdrawal (*p*=0.0140) timepoints compared to baseline. Moreover, EtOH-exposed uIntA rats drank significantly more than Air-exposed counterparts during both vapor (*p*<0.0001) and withdrawal (*p*=0.0016) timepoints. This is further supported by a significant main effect of vapor on change in intake from baseline observed during the vapor and withdrawal timepoints (**Fig 5C**). Maintenance of dependence-induced escalation was associated with a robust increase in eBECs. This was supported by a significant main effect of vapor and a significant vapor by timepoint interaction for within-session eBECs achieved during vapor exposure (**Fig. 5D**) and withdrawal (**Fig. 5E**). For both timepoints, Šidák’s posthoc comparisons did not reveal any simple main effects. A significant main effect of vapor and vapor by timepoint interaction was also observed in average peak eBEC during the final four sessions of each timepoint (**Fig. 5F**). Tukey’s posthoc comparisons indicate that peak eBEC was significantly higher in EtOH-exposed uIntA rats during the vapor (*p*=0.0207) and withdrawal (*p*=0.0150) timepoints compared to baseline. Furthermore, EtOH-exposed uIntA rats had higher peak eBECs compared to Air-exposed counterparts during vapor (*p*=0.0020) and withdrawal (*p*=0.0015) timepoints. Notably, aBECs measured at the end of the 24 hr withdrawal drinking session were not different between Air- and EtOH-exposed rats (t(16)=0.9582, *p*=0.3522, Air: 29.21±11.41 mg/dL, EtOH: 48.04±16.01 mg/dL). Nevertheless, a similar sustained elevation in bout probability was also observed in uIntA rats such that there was a main effect of vapor exposure on within-session bout probability for drinking sessions occurring during both vapor (**Fig. 5G, Supp Fig 2F,H**) and withdrawal (**Fig. 5H, Supp Fig 2G-H**) timepoints. Additionally, a main effect of vapor exposure was observed for bout probability AUC during the final four sessions of each timepoint (**Fig. 5I**). While not significant utilizing Fischer’s exact test, a trend for a higher proportion of rats defined as escalators (≥20% increase in intake from baseline) was apparent in EtOH- compared to Air-exposed rats during vapor exposure (**Fig 5J**; *p*=0.0567). This trend persisted albeit to a somewhat diminished degree, during withdrawal (**Fig 5K**; *p*=0.1534). Altogether, these data indicate that uIntA facilitates robust dependence-induced escalation of ethanol consumption that emerges during vapor exposure and is maintained at least four days into withdrawal.

**Figure 5:**
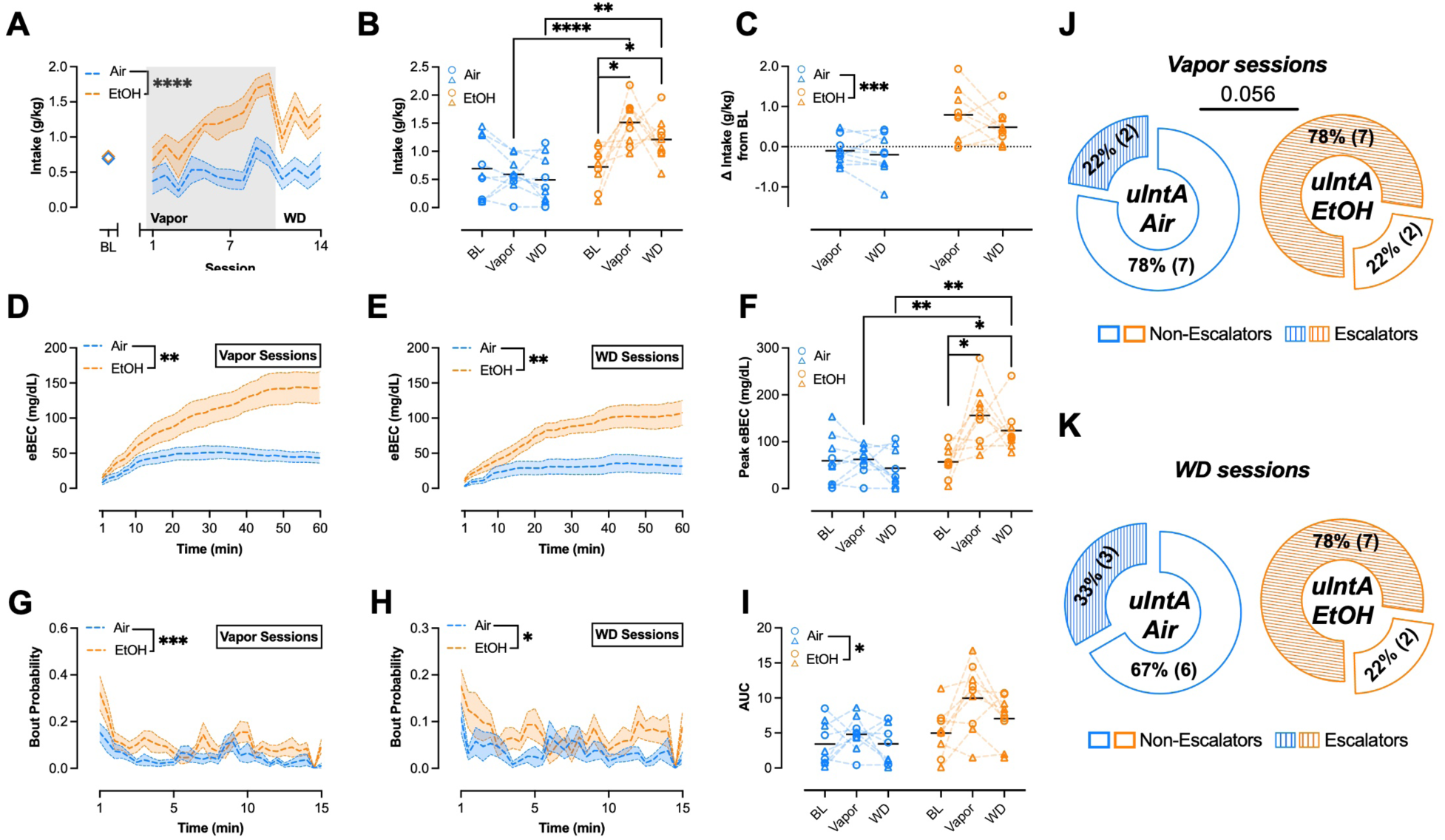
Long-lasting dependence-induced escalation of ethanol consumption in rats drinking under an ulntA schedule. **A)** Daily ethanol intake during vapor exposure and withdrawal timepoints relative to average intake at baseline. **(B)** Average intake across all timepoints. **(C)** Change in intake from baseline during vapor exposure and withdrawal. **(D)** Average within-session eBECs achieved during vapor exposure drinking sessions. **(E)** Average within-session eBECs achieved during withdrawal. **(F)** Average peak eBEC achieved across all timepoints **(G)** Average within-session bout probability during vapor exposure. **(H)** Average within-session bout probability during withdrawal. (I) Area under the curve for bout probability across all timepoints. Proportion of Air- and EtOH-exposed ulntA rats classified as escalators (≥ 20% increase from baseline) or non-escalators during vapor exposure **(J)** and withdrawal **(K)** sessions. Error bars indicate ± SEM. Dashed lines and open datapoints indicate ulntA. Circles and triangles indicate males and females, respectively. n=4-5/sex/group.

### Dependence-induced escalation of ethanol intake is greater in rats drinking on an uIntA schedule

Because we hypothesized *a priori*, that uIntA would facilitate greater dependence-induced escalation of ethanol intake than ContA, we performed an additional set of analyses examining the effect of schedule on dependent measures in only those rats that were exposed to ethanol vapor. This analysis detected a main effect of schedule on intake across drinking sessions (**Fig. 6A**) indicating that uIntA rats drank significantly more than ContA rats in all post-baseline drinking sessions. This is further corroborated by a significant main effect of schedule on intake averaged across the final four sessions at each timepoint (**Fig. 6B**). While not significant, there was a trending main effect of schedule (*p*=0.0773) on change in intake from baseline during vapor and withdrawal timepoints (**Fig. 6C**). A significant schedule by time interaction was detected in within-session eBEC dynamics observed during the final four vapor exposure drinking sessions (**Fig. 6D**). However, Šidák’s posthoc comparisons did not indicate any significant simple main effects. A significant main effect of schedule and schedule by time interaction were identified in within-session eBEC dynamics during withdrawal (**Fig. 6E**). Šidák’s posthoc comparisons found that uIntA rats achieved higher eBECs during minutes 29-32 of the drinking session compared to ContA rats. The ability of uIntA to promote higher levels of intoxication was further supported by a significant main effect of schedule on average peak eBECs across timepoints (**Fig. 6F**). In agreement, EtOH-exposed uIntA rats exhibited significantly higher aBECs at 24 hr withdrawal compared to EtOH-exposed ContA counterparts (t(17)=2.578, *p*=0.0195, ContA; 8.64±2.02 mg/dL, uIntA: 48.04±16.01 mg/dL). Similar results were observed with respect to bout probability with a significant schedule by time interaction found when within-session bout probability was assessed during the final four vapor drinking sessions (**Fig 6G).** Examination of the linear predictions of bout probability and their respective beta estimates at this timepoint suggests that EtOH-exposed uIntA animals exhibit a less robust decrease in bout probability over ethanol availability time compared to EtOH-exposed ContA animals (**Supp Fig 2I,K**). Moreover, a significant main effect of schedule was observed for within-session bout probability during withdrawal (**Fig. 6H, Supp Fig 2J-K**). This was further supported by a significant main effect of schedule on bout probability AUC assessed across all timepoints (**Fig. 6I**). Finally, Fischer’s exact test identified that the proportion of rats defined as escalators (≥20% increase in intake from baseline) was not significantly different between ContA and uIntA rats in drinking sessions that occurred during vapor exposure (**Fig 6J**; *p*>0.9999). However, a trending increase in the proportion of rats defined as escalators was observed in the uIntA group compared to the ContA group during withdrawal (**Fig 6K**; *p*=0.0698). Altogether, these data show that the uIntA schedule elicits a more robust and sustained dependence-induced escalation of ethanol consumption compared to the ContA schedule.

**Figure 6:**
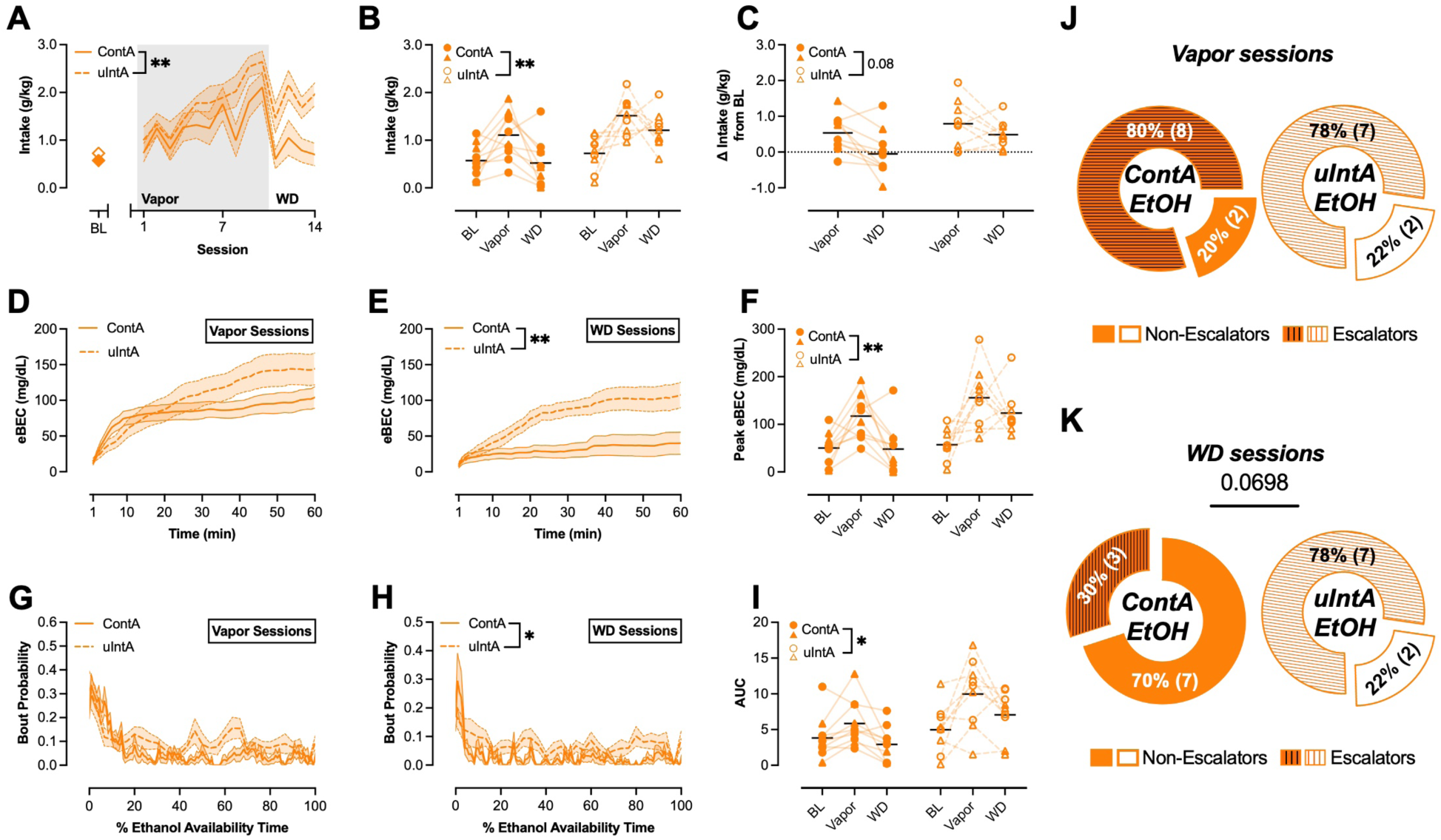
ulntA exacerbates dependence-induced escalation of ethanol consumption relative to ContA. **A)** Daily ethanol intake during vapor exposure and withdrawal timepoints relative to average intake at baseline. **(B)** Average intake across all timepoints. **(C)** Change in intake from baseline during vapor exposure and withdrawal. **(D)** Average within-session eBECs achieved during vapor exposure drinking sessions. **(E)** Average within-session eBECs achieved during withdrawal. **(F)** Average peak eBEC achieved across all timepoints **(G)** Average within-session bout probability during vapor exposure drinking sessions. **(H)** Average within-session bout probability during withdrawal. (I) Area under the curve for bout probability across all timepoints. Proportion of EtOH-exposed ContA and EtOH-exposed ulntA animals defined as escalators (≥ 20% increase from baseline) or non-escalators during vapor exposure **(J)** and withdrawal **(K)** sessions. Error bars indicate ± SEM. Solid lines and closed datapoints indicate ContA while dashed lines and open datapoints indicate ulntA. Circles and triangles indicate males and females, respectively. n=4-6/sex/group.

### uIntA affords additional opportunities to assess motivation and drinking strategies

Motivation to consume ethanol is often extrapolated by measuring effort to perform an operant response required to gain ethanol access. However, prior research has shown that an animal’s willingness to a perform an operant response is not necessarily linked to reinforcer consumption (Blegen et al., 2018; Patwell et al., 2021; Pitock et al., 2025; Puaud et al., 2018). Separately, a bevy of literature has demonstrated that motivation to consume liquid reinforcers can be extrapolated from lick microstructure (Dwyer, 2012; Johnson, 2018; Naneix et al., 2020; Spector, 2000). This work has linked parameters related to the frequency and timing of drinking bouts (i.e., latency to 1^st^ bout) with incentive to consume, whereas parameters related to bout duration (i.e., seconds; number of licks) and lick timing (i.e., interlick interval) with palatability of the reinforcer. In combination with the discrete ethanol access periods inherent to uIntA, these measures permit assessment of relative engagement during ethanol access periods that can be leveraged to evaluate motivation. With this in mind, we analyzed how drinking microstructure was affected by vapor exposure within uIntA rats.

As expected, the number of drinking bouts per DS presentation was significantly different between Air- and EtOH-exposed uIntA rats as evidenced by a significant main effect of vapor group (**Fig. 7A**). The percent of DS presentations where rats engaged (i.e., performed ≥2 bouts) was used as an indirect measure of bout frequency. This analysis revealed a significant vapor by timepoint interaction (**Fig. 7B**), with Tukey’s posthoc comparisons indicating that EtOH-exposed uIntA rats exhibit higher percent DS engagement in drinking sessions during vapor exposure compared to baseline (*p*=0.0498). Percent DS engagement was also greater in EtOH- than Air-exposed counterparts during both vapor (*p*=0.0003) and withdrawal (*p*=0.0390) timepoints. Furthermore, we observed a significant main effect of vapor on latency to first bout within each DS presentation (**Fig. 7C**). Due to the unpredictable nature of DS duration, we next considered the number of drinking bouts terminated by the end of a DS presentation, defined by performing the last lick of a drinking bout within 0.5s of DS offset (**Fig. 7D**). This analysis revealed a significant vapor by timepoint interaction with Tukey’s post hoc comparisons showing that EtOH-exposed uIntA rats exhibit more terminations during the vapor timepoint compared to baseline (*p*=0.0079). Moreover, EtOH-exposed uIntA rats exhibit more terminations than Air-exposed counterparts during both vapor (*p*=0.0004) and withdrawal (*p*=0.0454) timepoints. On the other hand, we observed a significant vapor by timepoint interaction on bout duration (**Fig. 7E**) with Tukey’s multiple comparisons indicating that bout duration was shorter in Air-exposed rats during withdrawal compared to baseline (*p*=0.0488). However, we did not observe an effect of vapor exposure or timepoint in bout length (**Fig. 7F**). Finally, we observed a significant vapor by timepoint interaction on interlick interval (**Fig. 7G**), with Tukey’s multiple comparisons tests indicating that Air-exposed rats exhibited a shorter interlick interval during the vapor timepoint than at baseline (*p*=0.0216). Altogether, these data suggest that dependence-induced escalation observed in EtOH-exposed animals is driven by increased incentive for ethanol, with the palatability of ethanol remaining stable in EtOH-exposed rats throughout the testing period.

**Figure 7:**
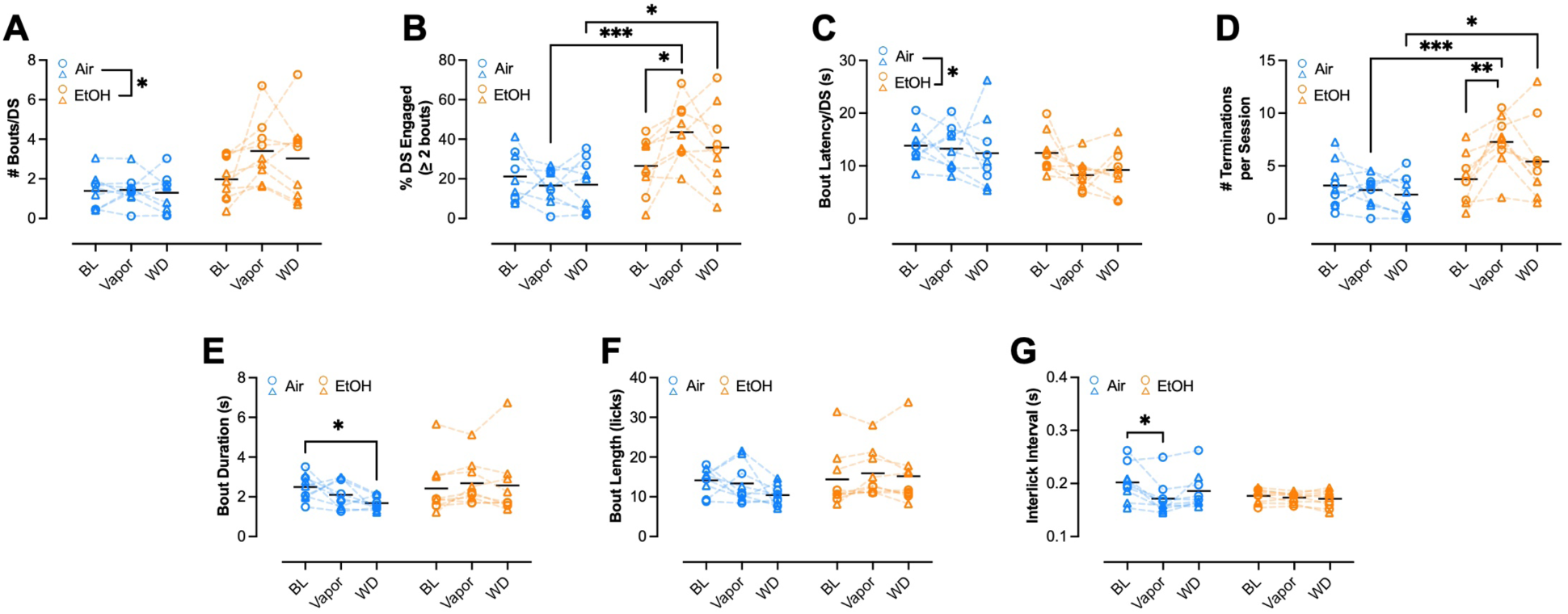
Ethanol dependence increases motivation to consume ethanol in rats drinking under an ulntA schedule. **(A)** Average number of bouts per OS presentation across all timepoints. **(B)** Percent of OS presentations engaged (≥ 2 bouts within the OS) across all timepoints. **(C)** Average bout latency from OS onset across all timepoints. **(D)** Average number of bouts terminated by OS offset across all timepoints. **(E)** Average bout duration across all timepoints. **(F)** Average bout length across all timepoints. **(G)** Average interlick interval across all timepoints. Circles and triangles indicate males and females, respectively. n=4-5/sex/group.

Due to the unpredictable nature of DS and ITI duration and timing within the uIntA schedule, we next performed a series of analyses to determine whether time within a given DS presentation, latency of DS onset (i.e. earlier vs. later in a session), bout termination by prior DS offset, duration of the prior ITI period, amount of ethanol consumed during the prior DS, and cumulative ethanol consumed affected bout probability during the current DS presentation. During baseline sessions (**Fig 8A-F, Supp Fig 3A-G**), there was an expected significant main effect of DS time on current DS bout probability (**Fig 8A, Supp Fig 3A**), such that bout probability increases across DS time. There was also a significant main effect of latency to DS onset on bout probability such that rats exhibited higher bout probability during early than late DS presentations (**Fig. 8B, Supp Fig 3B**). Moreover, a significant main effect of prior ITI duration was detected, such that rats exhibited higher bout probability following a longer duration ITI compared to shorter duration ITI (**Fig. 8C, Supp Fig 3C**). There was also a significant main effect of bout termination by prior DS offset on bout probability, with termination in the prior DS increasing bout probability during the current DS presentation period (**Fig. 8D, Supp Fig 3D**). This suggests that rats may be motivated to consume more in the subsequent DS when they are prematurely ‘cut-off’ from drinking in the prior DS. Furthermore, we also detected a significant main effect of the amount of ethanol consumed in the prior DS presentation period on current DS bout probability such that rats exhibited higher bout probability when they drank more in the prior DS compared to when they drank less (or not at all) (**Fig. 8E, Supp Fig 3E**). We also identified a significant main effect of cumulative ethanol consumption, such that bout probability in the current DS decreases as cumulative ethanol consumed increases (**Fig. 8F, Supp Fig 3F**). Examination of the beta estimates derived for each of these parameters reveals the relative effect of each term (**Supp Fig 3G**), indicating that experience of a termination in the prior DS and latency of DS onset served as the parameters most influential over current DS bout probability.

**Figure 8:**
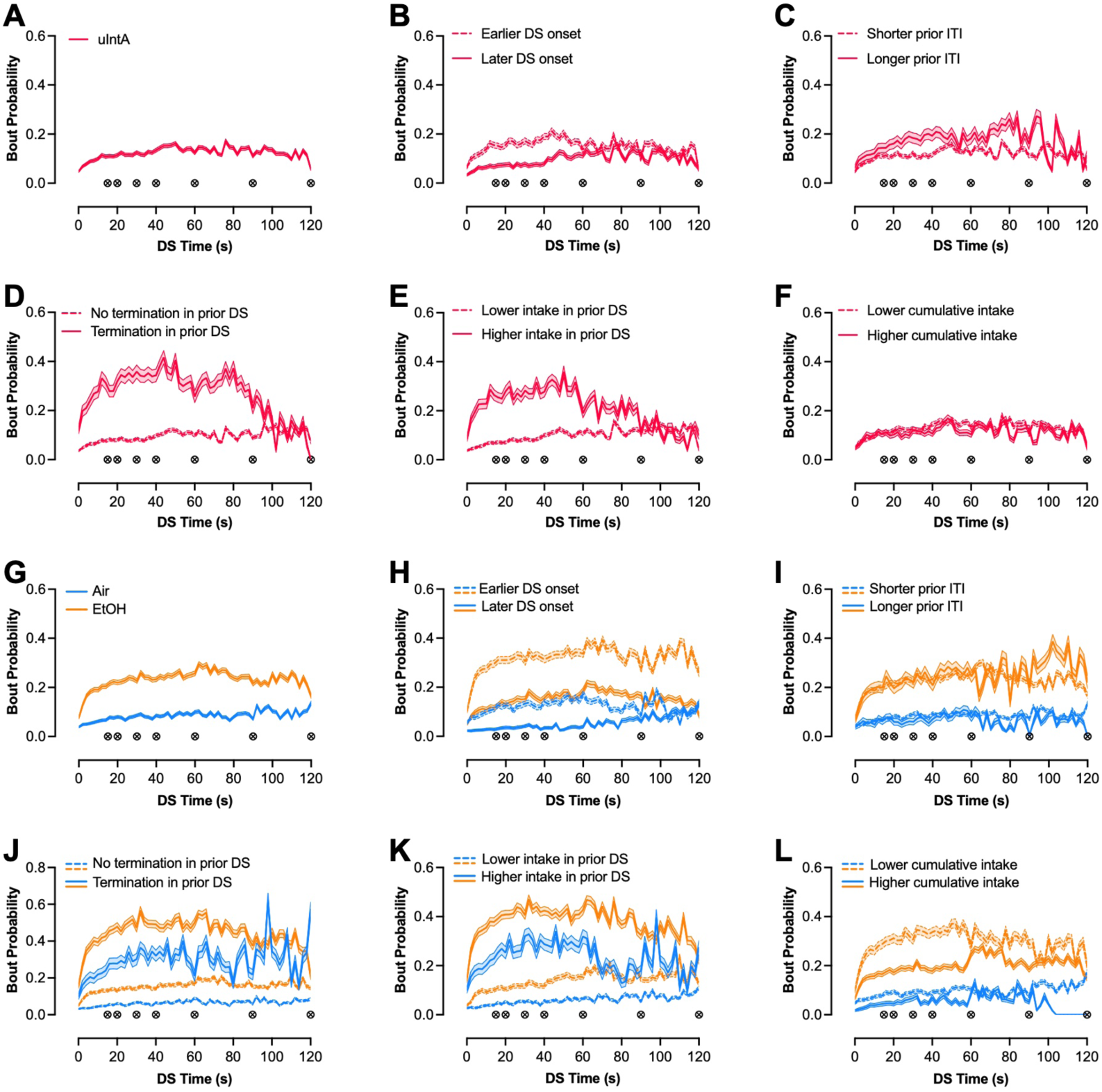
Factors affecting bout probability during DS presentations under the ulntA schedule of ethanol drinking. **(A)** Bout probability across DS presentation time for all DS presentations within the ulntA schedule during baseline drinking sessions. Effect of **(B)** latency of DS onset, **(C)** duration of ITI prior ITI period, **(D)** experience of a bout terminated at the end of the prior DS, **(E)** amount of EtOH consumed in the prior DS period, and **(F)** cumulative EtOH consumed on current DS bout probability during baseline drinking sessions. **(G)** Bout probability across DS presentation time for all DS presentations for Air-exposed and EtOH-exposed ulntA rats during drinking sessions occurring at the vapor and withdrawal timepoints. Effect of **(H)** latency of DS onset, (I) duration of ITI prior ITI period, **(J)** experience of a bout terminated at the end of the prior DS, **(K)** amount of EtOH consumed in the prior DS period, and **(L)** cumulative EtOH consumed on current DS bout probability across Air-exposed and EtOH-exposed rats during drinking sessions occurring at the vapor and withdrawal timepoints. Inset symbols indicate unpredictable offset of the DS period (i.e. 15, 20, 30, 40, 60, 90, or 120 s). *(n* = 4-5/sex/group)

Subsequent analyses were designed to determine whether ethanol dependence alters these drinking strategies by examining bout probability during the vapor and withdrawal timepoints (pooled data) (**Fig. 8G-L, Supp Fig 3H-N**). Importantly, the main effects observed for all independent variables assessed during the baseline sessions (**Fig 8A-F, Supp Fig 3A-G**) were retained at this timepoint, including DS time (**Fig. 8G, Supp Fig 3H**), latency of DS onset (**Fig. 8H, Supp Fig 3I**), prior ITI duration (**Fig. 8I, Supp Fig 3J**), experience of a termination in the prior DS (**Fig. 8J, Supp Fig 3K**), ethanol consumption in the prior DS (**Fig. 8K, Supp Fig 3L**), and cumulative ethanol consumption (**Fig. 8L, Supp Fig 3M**). In addition to these effects, significant interactions with vapor exposure were identified for DS time (**Fig 8G, Supp Fig 3H,N**), latency of DS onset (**Fig 8H, Supp Fig 3I, N**), experience of a termination in the prior DS presentation (**Fig 8J, Supp Fig 3K,N**), and ethanol consumption in the prior DS presentation (**Fig 8K, Supp Fig 3L,N**), such that EtOH-exposed animals exhibited a more robust influence of these variables on current DS bout probability compared to Air-exposed counterparts. Similar to the effect of these parameters during baseline sessions, examination of beta estimates derived for each of these parameters reveals that experience of a termination in the prior DS and latency of DS onset remained the most influential over current DS bout probability, with vapor exposure group exhibiting a similar magnitude of effect to these parameters (**Supp Fig 3N**). Moreover, these parameters were also identified to have the most robust interaction with vapor exposure (**Supp Fig 3N**). Altogether, these data suggest that ethanol dependence exacerbates front-loading of ethanol consumption, facilitates capitalization of the total DS duration, and may enhance drinking as a result of ‘frustration’ – factors that likely contribute to the emergence of escalated ethanol intake.

### uIntA promotes aversion-resistant drinking in ethanol-dependent rats

Finally, to determine whether uIntA promotes aversion-resistant drinking compared to ContA, we next assessed intake of ethanol adulterated with quinine (30 mg/L). There was a significant vapor by schedule interaction in intake observed during the quinine drinking session (**Fig. 9A**). Šidák’s multiple comparisons indicate that ethanol intake was higher in EtOH-exposed uIntA rats compared to Air-exposed uIntA (*p*=0.0316) and EtOH-exposed ContA (*p*=0.0066) rats. When intake was analyzed in terms of change from baseline, we observed a significant schedule by vapor interaction (**Fig. 9B**). Šidák’s multiple comparisons indicate that ethanol intake was significantly greater (i.e., more positive change) in EtOH-exposed uIntA rats compared to Air-exposed counterparts (*p*=0.0491) and EtOH-exposed ContA rats (*p*=0.0474). Surprisingly, despite EtOH-exposed uIntA rats achieving relatively higher aBECs by the end of the quinine-adulterated drinking session, this difference did not achieve statistical significance (**Fig. 9C**). However, a multifactorial RM ANOVA of eBEC dynamics during the quinine drinking session identified a significant three-way interaction between vapor, schedule, and time (**Supp Table 1**). As such, we performed separate two-way RM ANOVAs to assess the effect of vapor exposure on eBECs during the quinine drinking session. Within the ContA schedule, there was no effect of vapor exposure on eBECs (**Fig. 9D**). On the other hand, we observed a significant effect of vapor within the uIntA rats (**Fig. 9E**). This is corroborated by a significant vapor by schedule interaction in peak eBECs achieved during this session (**Fig. 9F**). Šidák’s multiple comparisons indicate that EtOH-exposed uIntA rats achieved a significantly higher peak eBEC than Air-exposed uIntA (*p*=0.0218) and EtOH-exposed ContA (*p*=0.0046) rats. Finally, we performed Fisher’s exact tests to compare the proportion of rats classified as aversion-resistant (≤20% decrease in intake from baseline) between groups. Two outliers were identified using ROUT outlier detection (*n* = 1 Air/ContA, *n* = 1 Air/uIntA) and were excluded from this analysis (**Fig. 9G**). We found a greater proportion of aversion-resistant rats in the EtOH-exposed uIntA group compared to Air-exposed counterparts (*p*=0.0034) utilizing Fischer’s exact test. The proportion of rats defined as aversion-resistant was also significantly greater in EtOH-exposed uIntA rats compared to EtOH-exposed ContA rats (*p*=0.0198) (**Fig 9G**). Importantly, there were no significant differences in the proportion of aversion-resistant vs -sensitive rats between ContA Air-exposed and EtOH-exposed groups (*p*>0.9999) or between the ContA Air-exposed and uIntA Air-exposed groups (*p*>0.9999). Altogether, these data suggest that uIntA promotes aversion-resistant drinking exclusively in ethanol dependent rats, unlike the ContA schedule.

**Figure 9:**
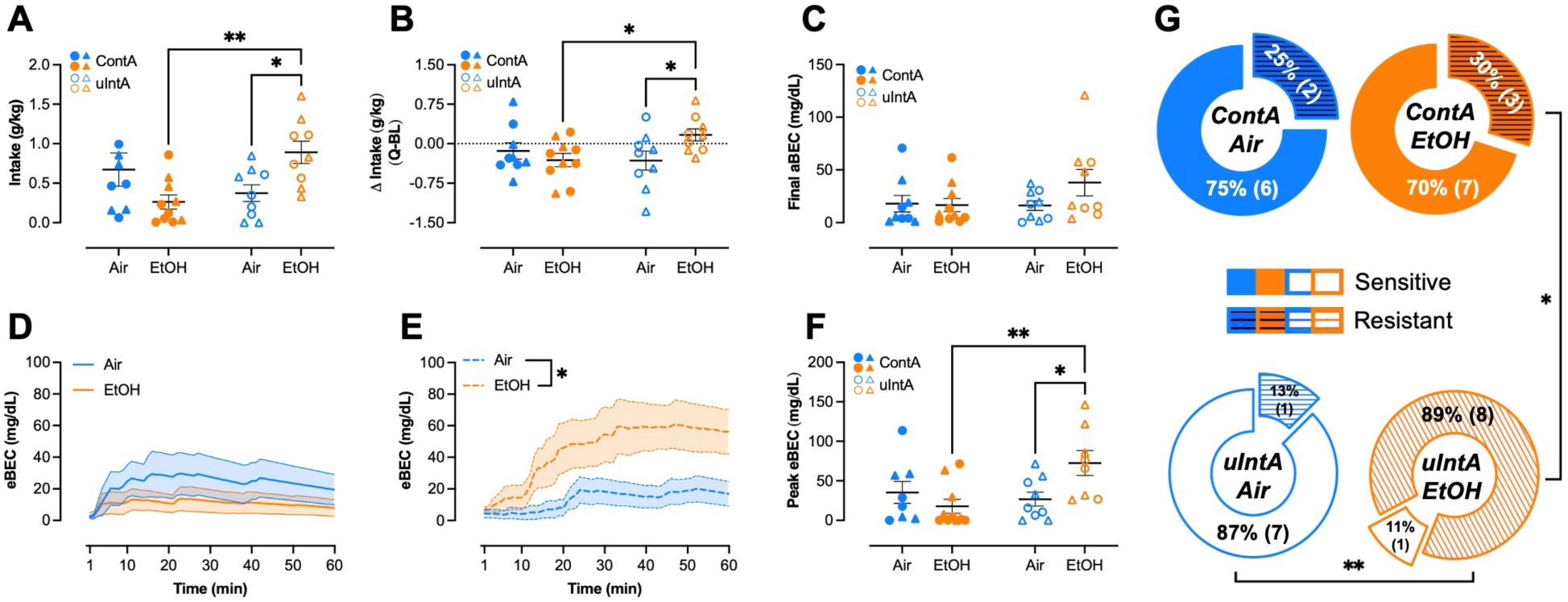
ulntA promotes dependence-induced aversion-resistant drinking. **(A)** Intake of quinine-adulterated ethanol on the final day of testing. **(B)** Change in intake of quinine-adulterated ethanol from average baseline (BL) intake. **(C)** End-of-session aBEC following quinine-adulterated ethanol consumption. eBEC for ContA **(D)** and ulntA **(E)** rats across vapor groups throughout the 60-min quinine-adulterated drinking session. **(F)** Peak eBEC achieved during the quinine-adulterated drinking session. **(G)** Proportion of rats within each group classified as aversion-sensitive or -resistant (≤ 20% decrease from baseline). Error bars indicate ± SEM. Solid lines and closed datapoints indicate ContA and dashed lines and open datapoints indicate ulntA. Circles indicate males and triangles indicate females. n = 4-6/sex/group.

## Discussion

Here, we present a novel uIntA schedule of voluntary ethanol intake that engenders similar levels of drinking prior to ethanol dependence to a ContA procedure, despite significantly shorter total duration of ethanol access (15 min vs. 60 min, respectively). uIntA exacerbates dependence- induced escalation and aversion-resistant drinking compared to a ContA procedure, suggesting that it may serve as a more suitable model of AUD-relevant symptomatology. Analysis of lick microstructure as well as parameters associated with the discrete ethanol access periods inherent to the uIntA paradigm provide additional measures related to the motivation to consume ethanol and variable within-session drinking patterns without the need for an operant response requirement. Importantly, this is achieved in the uIntA procedure with no greater experimental burden than the ContA procedure. Altogether, this work highlights uIntA as a powerful tool, particularly when used in combination with estimated BEC analyses, to elucidate pharmacological, psychological, and neurobiological factors contributing to loss of control over drinking within AUD.

While eBECs are used extensively in clinical studies of voluntary ethanol consumption (Hultgren et al., 2025; Piasecki, 2019), their use is not widespread in preclinical work. To our knowledge, the present study is the first to provide within-session estimates of BEC that can be used to support claims related to drinking patterns and overall ethanol intake. Variability in drug pharmacokinetics can dynamically modify intake, motivated responding, and SUD-related symptomatology across drug classes (Allain et al., 2015; Kawa et al., 2019; Spear, 2020; Zimmer et al., 2011). Thus, estimation of within-session BECs achieved by rodents during voluntary drinking sessions provides a critical translational stepping stone that enables researchers to evaluate the applicability of reported levels of intake to clinical manifestations of AUD, as was recently emphasized by Grahame (2026). For the current analyses, we utilized a modified application of the commonly described Widmark formula paired with a zero-order kinetic model of ethanol elimination (Widmark, 1981), allowing for accumulation of ethanol consumption across the session. We validated our eBEC pipeline using aBECs captured during voluntary ethanol consumption on three separate occasions throughout the experiment, as well as pharmacokinetic data previously established in Long-Evans rats (B. Webb et al., 2002). Individual estimation of the volume distribution of ethanol (*V_d_*) and barrier terms (*B*) within the range of values provided by the literature allowed us to predict end-of-session aBECs in outbred Long-Evans rats with strong fidelity (ave residual: +2 mg/dL, range: -6-+16 mg/dL). Nevertheless, there are some limitations to the current approach that should be noted.

Application of the Widmark equation in medicolegal contexts assumes (1) all ethanol has been consumed, (2) this ethanol has been fully absorbed, and (3) is a sufficient amount to saturate ethanol metabolic processes, isolating any changes in BEC to the zero-order, or linear, portion of hepatic metabolism (Jones, 2019; Posey & Mozayani, 2007). This is an important consideration to ensure accurate back-calculation of BEC at the time of an event with an error of ± 20 mg/dL (Gullberg, 2007; Maskell et al., 2017; Searle, 2015). In our modified application of this equation, we describe near-instantaneous absorption of ethanol in boluses defined by our binning window (1 min) that is limited by a barrier term (*B*) reflecting any process that could buffer ethanol availability in the blood (Goldman et al., 2025; Jones, 2019; Wilson & Matschinsky, 2020). As such, our eBEC estimates are limited by an incomplete account of the non-linear, volume-dependent dynamics of the gastric emptying rate that governs ethanol absorption, as well as a lack of separation of metabolic processes that could be differentially influenced by the volume and/or concentration of ethanol observed in a given compartment (Jones, 2019). Moreover, there are several interconnected factors that can affect the absorption, distribution, and metabolism of ethanol within a given individual across time, including age, sex, weight, current hydration, nutrition, stomach contents, fat distribution, genetics, pattern of drinking, and alcohol history (Cederbaum, 2012; Glover et al., 2021; Goldman et al., 2025; Jones, 2019). Furthermore, our analysis does not explicitly account for metabolic tolerance (affecting either first-pass metabolism in the gastric or hepatic compartments or systemic metabolism in the hepatic compartment) that may have developed over the course of ethanol vapor exposure. Individual estimation of *V_d_* and *B* values derived from multiple timepoints throughout testing controls for variation likely exhibited in these factors that we did not and could not measure both between and within animals. As such, the individually estimated *B* term in this context most likely reflects an amalgamation of individual differences exhibited in ethanol absorption and metabolism observed in orally self-administering rats that could be affected by the aforementioned factors. This includes variable absorption as a result of gastric concentration/volume, gastric metabolism prior to absorption, first-pass metabolism in the liver, and then hepatic ethanol metabolism from the systemic circulation (Goldman et al., 2025; Jones, 2019; Wilson & Matschinsky, 2020). It is important to note that while many studies suggest that chronic intermittent ethanol vapor exposure produces metabolic tolerance, no studies have quantified this systematically in Long-Evans rats and sparse work conducted in Wistar and Sprague Dawley rats as well as C57BL6/J mice has not identified significant differences in ethanol elimination rate following vapor exposure (Gilpin et al., 2009; Jury et al., 2017; Lopez et al., 2012; Rimondini et al., 2008). However, this lack of metabolic tolerance is concomitant with functional tolerance to the sedative properties of ethanol in rat strains (Rimondini et al., 2008; Rogers et al., 1979), suggesting that tolerance produced by vapor exposure may be due to altered pharmacodynamics rather than pharmacokinetics. Additional work in alcohol-preferring rodent strains (cHAP mice and P rats) showing heightened metabolic tolerance as a result of chronic ethanol exposure suggests that pharmacokinetic alterations may be unique to these models (T. K. Li et al., 1979; T. K. Li, 2000; Lumeng & Li, 1986; Matson et al., 2013). While it is well accepted that chronic alcohol use in clinical populations can evoke significant metabolic tolerance (Goldman et al., 2025), work is needed to systematically define the emergence and persistence of metabolic tolerance following chronic intermittent ethanol vapor exposure in outbred rodent strains to allow for consideration of this in future pharmacokinetic modeling. Future work should also seek to develop multi-compartment models for preclinical studies that can consider variable rates of gastric absorption, first-pass metabolism, uneven distribution into bodily compartments, and the non-linear components of gastric and hepatic ethanol metabolism. Moreover, much of the current literature has disproportionately focused on estimating ethanol pharmacokinetic parameters relevant to these models in male rodents, delineating a need to investigate females to provide global, sex-specific estimations of relevant parameters for development of physiologically-based pharmacokinetic models. Altogether, these limitations emphasize that the within-session eBECs calculated using the current approach are estimates that are unable to account for all possible variables influencing absorption, distribution, and metabolism and must be considered in tandem with aBEC measurements.

The aforementioned caveats aside, the findings from the present study underscore the utility of eBECs in preclinical drinking studies. Within-session time course data reveal the approximate time point at which subjects reach peak intoxication across each phase of the study. These estimates provide further evidence for front-loading and often capture maximal intoxication that is missed when blood samples are obtained at the end of a drinking session. This is readily apparent in the discrepancy between peak eBECs and end-of-session aBECs as a result of voluntary ethanol drinking at 24 hr withdrawal and the quinine drinking session. Moreover, using eBECs, we also show that while non-dependent rats maintained peak eBECs of ∼60 mg/dL, ethanol-dependent rats were able to achieve eBECs above the criteria for binge drinking ((NIAAA, 2025), >80 mg/dL). In some cases, peak eBECs exceeded this criterion reaching values >150 mg/dL. Importantly, previous work in Long-Evans rats report a similar range of aBECs after ethanol intake of similar or greater magnitude during 30 min voluntary drinking or operant self-administration sessions (Carnicella et al., 2009; Chappell et al., 2013; J. Li et al., 2011; Morales et al., 2015; Ortelli & Weiner, 2025; Simms et al., 2008, 2010) further supporting the fidelity of our intake measurements and eBEC analysis. Our data also show that ethanol-dependent uIntA rats achieved significantly higher eBECs than ethanol-dependent ContA rats during and after vapor exposure despite having a shorter duration of ethanol access. These results confirm the clinical relevance of the drinking and intoxication levels captured using the uIntA paradigm. Widespread use of within-session eBECs in future preclinical studies will provide the field with important perspective on the translational relevance of the model(s) used and outcomes observed (Grahame, 2026), which can provide valuable information regarding therapeutic interventions for AUD.

Prior work has shown that within-session IntA can elicit higher ethanol intake compared to ContA (Gage et al., 2023; Tomie et al., 2006). Additional work found that unpredictability of non-drug reinforcer access via uIntA further enhances reward seeking compared to IntA and ContA schedules (M. J. F. Robinson et al., 2023), prompting us to assess this schedule in the context of ethanol consumption. As such, it was surprising that we did not observe greater ethanol intake in uIntA compared to ContA rats at baseline in the present study. Of note, in Gage et al. (2023) rats were trained to lever press for ethanol under a ContA schedule for four weeks in 24 hr drinking sessions after which they transitioned to an IntA schedule where access to ethanol was progressively reduced from 40 to 2 min per hr. Although Tomie et al. (2006) did not require an operant response in their drinking paradigm, ethanol access in this study was limited to only 10 s at a time during a 30 min drinking session, with ethanol concentration increasing over time. As such, it is possible that simple methodological differences play a role in these discordant results. Furthermore, while (Gage et al., 2023) found that IntA increased seeking for ethanol (i.e., lever pressing) in male rats, this was not associated with a concurrent increase in ethanol consumption. Other work showing enhanced seeking for non-drug reinforcers using an IntA/uIntA schedule compared to a ContA schedule did not directly measure reinforcer consumption (Beasley et al., 2022; Muñoz-Escobar et al., 2019; M. J. F. Robinson et al., 2023). Additionally, prior work has shown that seeking does not directly predict consumption (Blegen et al., 2018; Patwell et al., 2021; Pitock et al., 2025; Puaud et al., 2018; Renteria et al., 2020). With this in mind, while we are unable to capture seeking in the current experimental design, it is possible that uIntA promotes heightened motivation to obtain ethanol (i.e. seeking) in non-dependent rats without altering consumption as compared to ContA. Indeed, we found that uIntA rats drink to a similar degree to ContA rats prior to ethanol dependence despite significantly less time with access to ethanol. This suggests that uIntA rats, who have to adapt their drinking patterns to these unpredictable ethanol access periods, may be more motivated to obtain ethanol prior to ethanol dependence. Future work could incorporate an operant requirement to the uIntA schedule developed in the present study to discriminate the effect of schedule on lever pressing versus drinking in this paradigm.

Our findings show that uIntA exacerbates dependence-induced escalation of ethanol consumption relative to ContA. Interestingly, while ContA rats exhibited escalated intake in parallel with vapor exposure, the magnitude of escalation was lower than uIntA rats and transient, returning to baseline levels once vapor exposure ended. In contrast, escalated intake was sustained into acute withdrawal in uIntA rats. Foundational work using IntA theorized that ‘spiking’ drug concentrations produced by intermittent reinforcer intake drives SUD-related phenotypes for psychostimulants (Allain & Samaha, 2019; Kawa et al., 2019; Zimmer et al., 2011, 2012). However, recent work has identified that IntA/uIntA for non-drug reinforcers can drive similar phenotypes (Beasley et al., 2022; Muñoz-Escobar et al., 2019; M. J. F. Robinson et al., 2023), suggesting that the temporal nature of consumption, and/or resultant ‘spiking’ drug concentrations, are not necessarily required for this effect. On the other hand, enhanced seeking resulting from intermittent reward access may be due to processes related to incentive salience, defined as the motivational property of ‘wanting’ a reinforcer, which is separable from ‘liking’ (Olney et al., 2018). Inherent to IntA and uIntA paradigms, rats are presented with numerous opportunities to pair a neutral stimulus with reinforcer availability, likely resulting in a robust association between DS presentation and reinforcer access. As such, the strength of this association may imbue the DS with greater motivational salience in uIntA rats compared to ContA. Importantly, compared to ContA, the unpredictable and intermittent nature of DS presentations inherent to uIntA more closely reflects the variable cue dynamics experienced by humans (i.e. friends, music, physical setting, etc.), which are paired with and modulate drinking (Dvorak et al., 2014; Fischer et al., 2023; Fridberg et al., 2021; King et al., 2025; Pelloux et al., 2019; Prignitz et al., 2024; Wray et al., 2014). Like in uIntA, these cues are often repeatedly paired with ethanol availability, and may, over an extended alcohol history, be sufficient to drive motivated seeking for ethanol (Cofresí et al., 2019). As such, repeated stimulus pairings, and willingness to engage/consume during these pairings in the context of ethanol dependence, may elicit incentive sensitization (T. E. Robinson & Berridge, 2025). In studies using highly potent reinforcers like cocaine and opioids, the neurobiological processes underlying incentive sensitization may be more easily engaged, thereby allowing IntA to drive SUD-related phenotypes in the absence of physical dependence (Allain & Samaha, 2019; Beasley et al., 2023; Ramborger et al., 2026). In contrast, in the context of ethanol, a less rapidly potent reinforcer relative to other addictive drugs, we found that the ability of uIntA to drive escalation is only apparent in ethanol-dependent rats. These data suggest that uIntA alone is not sufficient to drive incentive sensitization for ethanol-associated cues but, rather, capitalizes upon dependence-induced neurobiological plasticity (Cofresí et al., 2019), leading to an exacerbation of dependence-induced escalation. Importantly, the present study is not able to disambiguate the effects of incentive salience to the DS and the discrete, temporal nature of drug intake in promoting this phenotype. Similarly, whether the unpredictable nature of DS presentations acts to augment the observed effects over predictable IntA, as reported previously for non-drug reinforcers (M. J. F. Robinson et al., 2023), is unknown. Future work comparing a uIntA schedule with and without a DS to IntA can help to address these gaps.

Critically, it is possible that metabolic and/or functional tolerance contributed to the escalated ethanol intake observed in ethanol-dependent rats in the current study. However, as mentioned earlier, evidence for the development of metabolic tolerance following CIE vapor exposure is scant. In addition, it is important to note that rats in the ContA and uIntA groups underwent the same ethanol vapor exposure and therefore should develop similar levels of tolerance. Thus, while it is possible that a portion of the escalation observed in both ContA and uIntA rats during ethanol vapor exposure is aimed at overcoming acute tolerance (Comley & Dry, 2020; Holland & Ferner, 2017), the escalation maintained into withdrawal in uIntA rats is likely driven by a separate motivation, such as heightened incentive salience to the DS. Indeed, clinical work has shown that acute metabolic tolerance produced by heavy alcohol exposure can be reversed within 1-3 days of abstinence (Keiding et al., 1983). As mentioned previously, additional work systematically defining the development of metabolic tolerance as a result of vapor exposure is required to better appreciate the role that this would play in escalation of ethanol consumption.

Prior literature has identified that lick microstructural analysis can extrapolate measures of incentive value and reinforcer palatability (Dwyer, 2012; Johnson, 2018; Naneix et al., 2020; Renteria et al., 2020; Spector, 2000). Using lick microstructural analyses, we found that EtOH- exposed uIntA rats had more bouts per DS and decreased latency to first drinking bout with no change in bout duration, bout length, and interlick interval. This suggests that dependence-induced escalation of ethanol consumption observed in uIntA rats is due to a change in the incentive value of ethanol as a reinforcer as opposed to a change in the palatability of ethanol. While not examined in the current study due to the configuration of our lickometer-equipped apparatus, assessment of changes in lick duration could provide more evidence toward the emergence of drinking patterns that seek to maximize ethanol intake as a result of dependence, especially in light of our eBEC analysis (Mackowiak et al., 2025; Petersen et al., 2024; Renteria et al., 2020). In addition to these measures, compared to AIR-exposed counterparts, EtOH-exposed uIntA rats had greater DS engagement and a higher number of terminated drinking bouts per session concomitant with dependence-induced escalation. Assessing the degree to which a rat engages (or not) when presented with ethanol access has the potential to serve as a translationally relevant outcome associated with an individual’s perceived loss of control over consumption when alcohol is freely available (Leeman et al., 2009, 2010; Patock-Peckham & Corbin, 2023; Remmerswaal et al., 2019; Waddell et al., 2025). This measure may also provide an avenue to further assess the motivational potency of an ethanol-associated cue to drive consumption (Sayette, 2016; Vafaie & Kober, 2022). On the other hand, terminated drinking bouts further elaborates upon the intensity of drinking occurring within each DS presentation similar to latency to first bout. Importantly, the underlying basis for increased motivation captured by these measures remains unclear and could be driven by any number or combination of factors, including but not limited to negative reinforcement (i.e. heightened drinking to alleviate withdrawal symptoms), increased incentive salience to the DS (as discussed), and altered sensitivity to the rewarding and/or aversive properties of ethanol consumption.

Unpredictable access periods inherent to uIntA have the potential to promote drinking strategies that seek to maximize ethanol intake within a limited amount of time (i.e. binge) (Jeanblanc et al., 2019). This may be driven, at least in part, by the perception of scarcity of ethanol access as a result of these unpredictable periods. Intuitively, adaptive processes in mammals seek to maximize the reinforcing value of food and motivation for food-seeking in uncertain conditions (Anselme & Güntürkün, 2019; Crandall & Temple, 2018; J. Webb et al., 2025), suggesting that introduction of unpredictability for drug reinforcers may impinge upon these adaptive processes to drive maladaptive intake. As such, we were particularly interested in quantifying a maladaptive front-loading consumption phenotype, which is thought to be driven by and enhance the rewarding effects of intoxication (Ardinger et al., 2022). However, criteria to define front-loading and its analysis in preclinical assessments are often ambiguous and variable (Ardinger et al., 2022), delineating a need in the field to better identify and isolate this phenotype. Utilizing linear mixed effect modeling, we found that DS onset latency was one of the most robust predictors of current bout probability, with uIntA rats exhibiting higher bout probability at earlier DS presentations. Additional moderate effects of ethanol consumption in the prior DS and time within the DS were also identified, with uIntA rats exhibiting higher bout probability when they drank larger quantities of ethanol in the preceding DS presentation and later within DS presentations. These parameters point to a significant concentration of drinking in earlier portions of the session, highlighting a front-loading strategy that seeks to maximize the rewarding properties of ethanol in non-dependent rats. We also identified a moderate effect of prior ITI duration, such that non-dependent rats exhibited higher bout probability in DS presentations preceded by a longer ITI duration. Surprisingly, experience of a bout terminated by the prior DS offset served as the most robust predictor of current DS bout probability. These effects may point to the development of perceived scarcity in ethanol availability time, as well as frustration associated with being ‘cut off’ from drinking. Finally, we observed a moderate effect of cumulative ethanol consumption, such that current bout probability decreased as cumulative ethanol consumption increased throughout the session. Interestingly, with the exception of ITI duration and cumulative ethanol consumption, each of the factors we found to alter bout probability were significantly exacerbated in ethanol-dependent rats at the time when dependence-induced escalation was observed. Together, these results suggest that parameters associated with uIntA elicit behavioral strategies that seek to maximize ethanol intake and are leveraged in ethanol-dependent rats to exacerbate dependence-induced escalation. Further, this work highlights a functionally relevant front-loading phenotype that is heightened by ethanol dependence, providing an avenue to rigorously examine the relevance of this phenotype to clinical manifestations of AUD. Future work should seek to incorporate comparison to an IntA schedule to better isolate the effect of uncertainty associated with DS presentations and identify how it contributes to these drinking phenotypes.

Finally, we found that ethanol-dependent uIntA rats exhibited significant insensitivity to quinine-adulterated ethanol compared to non-dependent counterparts and all ContA rats. Crucially, however, sensitivity to quinine was not particularly high in non-dependent or ContA rats. While previous studies have shown sensitivity to the same concentration of quinine used in the current study (Hopf et al., 2010; Katner et al., 2022; Lesscher et al., 2010; Loi et al., 2010; Nentwig et al., 2019), testing with multiple increasing concentrations of quinine may provide more conclusive evidence of the development of dependence-induced aversion resistance in uIntA rats by allowing for comparison of drinking at a concentration that produced greater sensitivity in non-dependent animals. However, visual inspection of the data (**Fig 9B**) reveals substantial individual variability in the ability of quinine adulteration to decrease ethanol intake in non-dependent uIntA and ContA rats. This points to the potential for individual differences in aversion-resistance independent of ethanol dependence. In agreement with this, prior work has found significant individual variability in aversion-resistant drinking and has shown that this phenotype preceded chronic ethanol exposure as opposed to developing as a result of ethanol dependence (Augier et al., 2018; Bluitt et al., 2025). This notion is further supported by clinical research suggesting that facets of impulsivity relating to risk tolerance are associated with vulnerability to future heavy drinking and AUD diagnosis (Fernández-Artamendi et al., 2018; Stamates & Lau-Barraco, 2017) and may also be exacerbated by heavy drinking/AUD diagnosis (Gerst et al., 2017; Rossiter et al., 2012; Sistad et al., 2019; Takahashi et al., 2009). Thus, the relatively small differences in aversion-resistant drinking observed in the current study may reflect a combination of innate individual differences as well as consequences of drinking and chronic ethanol vapor exposure. Future work using a within-subjects design to characterize aversion resistant drinking before and after dependence can help to address this outstanding question.

In conclusion, our newly developed uIntA ethanol drinking paradigm more faithfully models the often unpredictable nature of access to alcohol experienced by humans, promotes maladaptive drinking patterns, and significantly exacerbates dependence-induced escalation and aversion-resistant ethanol consumption. In combination with estimated measures of intoxication, this model represents a powerful tool that can be used to further our understanding of psychological and neurobiological factors underlying loss of control over drinking.

## Acknowledgements

The authors thank Nikki Kinarasri and Shikun Hou for technical support. This work was supported by NIH grants R01 AA031003, R01 AA029130, and P50 AA922537 to EJG and T32 AA0236577 to EHM.

---

**Supplementary Figure 1:**
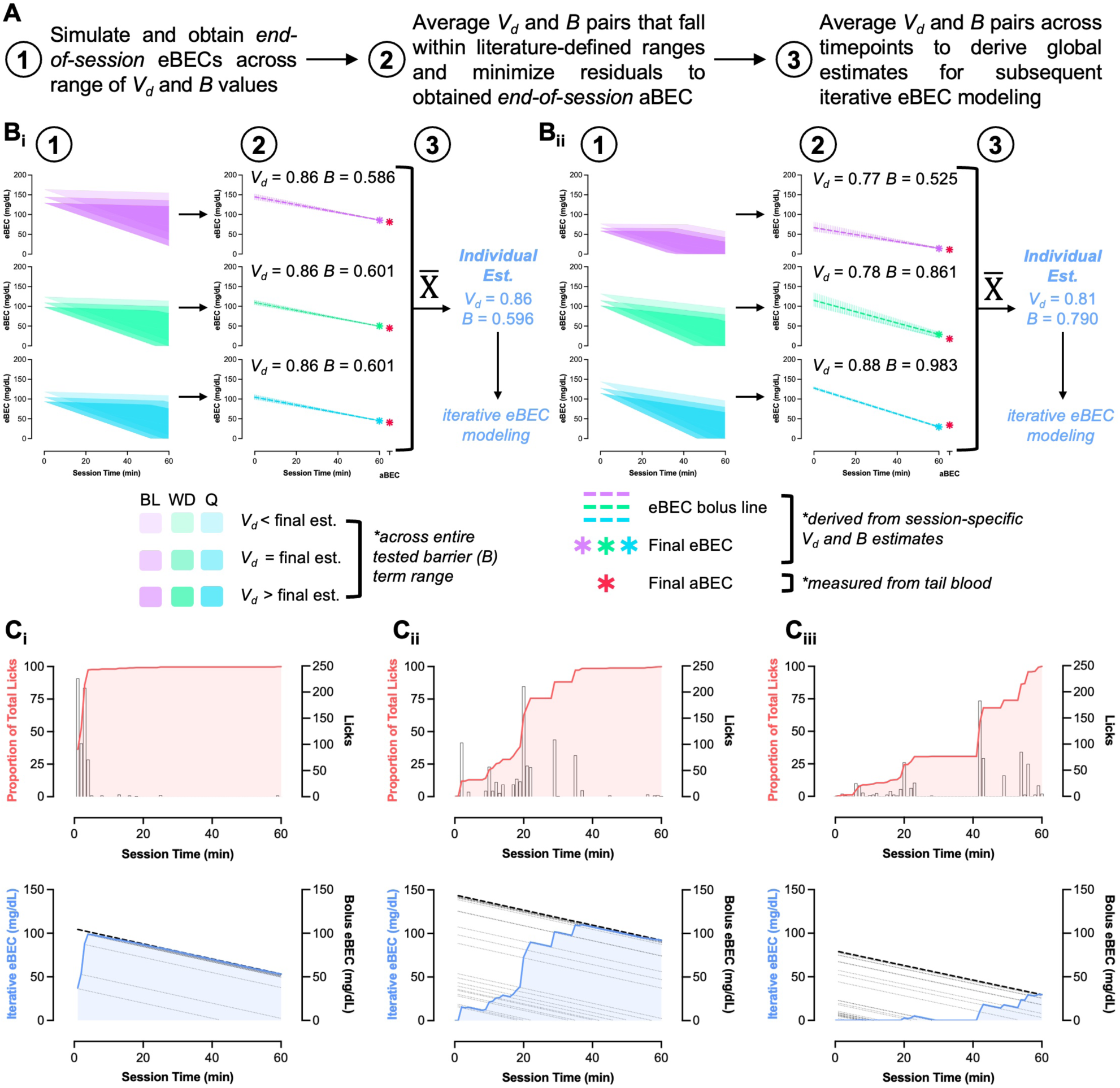
Individual estimation of volume of distribution (V_d_) and barrier (B) terms and lickometry-paired eBEC modeling. **(A)** Description of process to derive individual estimations of *V_d_* and *B* terms for iterative eBEC modeling. **(B_i-ii_)** Representative visualizations of processes described in A. Firstly, ‘end-of-session’ eBEC calculations for sessions where tail blood was collected are derived from the range of *V_d_* and *B* terms tested. Importantly, this is modeled as if the animal’s total intake occurred at session start, similar (but not identical) to how the Widmark equation would be applied in clinical and medicolegal settings. Secondly, the *V_d_* and *B* term pairs that produce ‘end-of-session’ eBEC calculations that minimize residuals compared to the obtained ‘end-of-session’ aBEC value and fall within literature-defined ranges are averaged within each of these sessions. Thirdly, session-specific values are averaged to arrive at global *V_d_* and *B* estimates for each animal that are then used to calculate eBECs iteratively for all experimental sessions. Of note, Bii provides an example of how differences in these estimates may be identified throughout our experimental timeline, allowing for control of possible intraindividual variability produced by extraneous factors over time. **(C)** Visualization of iterative eBEC modeling upon derivation of global *V_d_* and *B* constants in which the animal reached their total intake early in the session **(C_i_),** in the middle of the session **(C_ii_),** and at the end of the session **(C_m_)-** Importantly, iterative eBEC modeling (blue line, bottom) does not match the ‘bolus’ eBEC line (dashed black line, bottom) derived from modeling all intake occurring at session start until the animal reaches their total intake for the session (red, top), limiting the achievable BEC scaled by the animal’s total intake as well as controlling for the rate of intake across the session

**Supplementary Figure 2:**
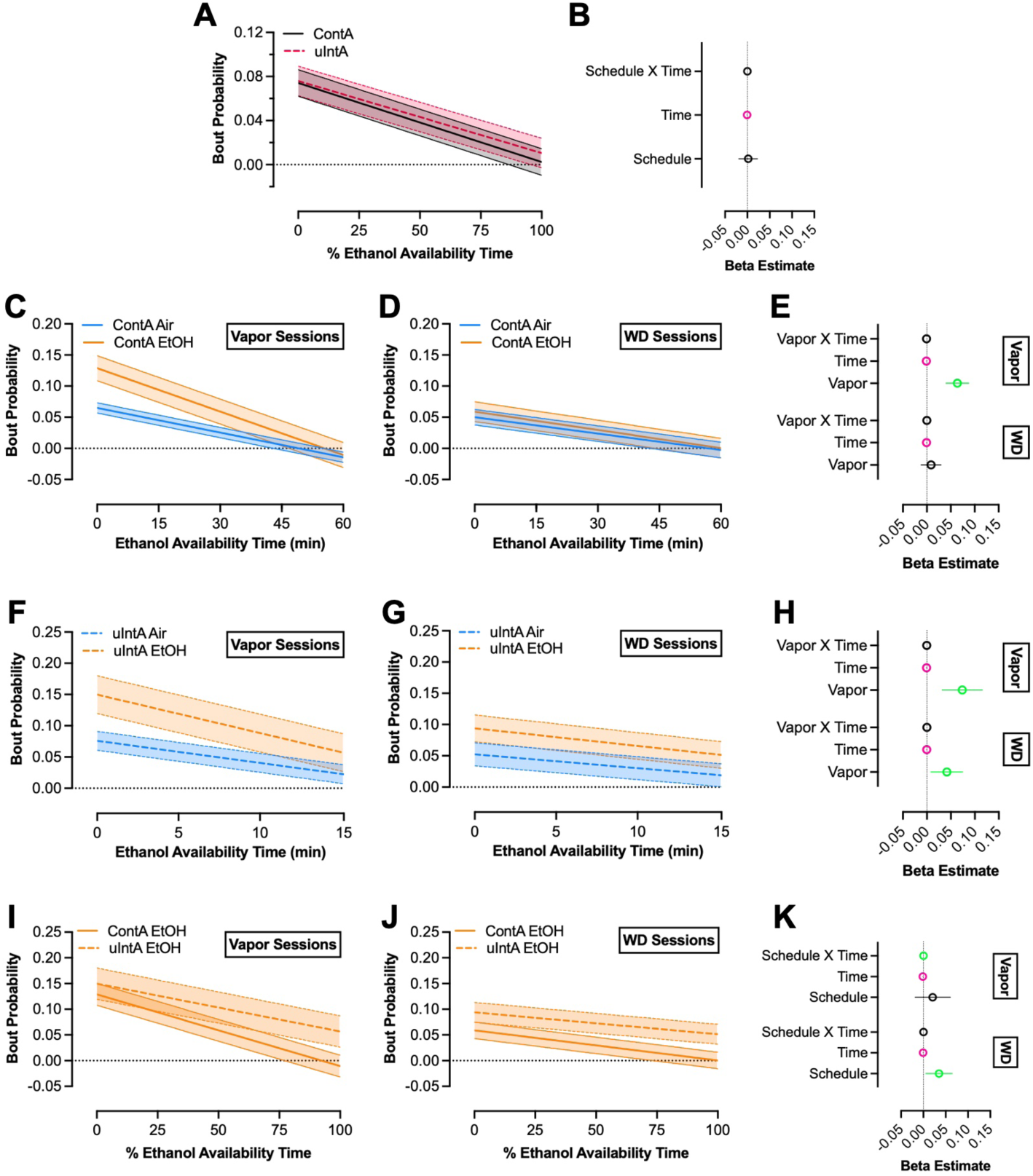
Linear predictions and beta estimates for the effect of vapor and schedule on bout probability across ethanol availability time. Linear prediction of bout probability between ContA and ulntA **(A)** and beta estimates **(B)** during the final four sessions of baseline drinking derived from data presented in Figure 3. Linear prediction of bout probability in ContA Air- and EtOH-exposed animals for the final four sessions of vapor exposure **(C)** and withdrawal **(D),** with beta estimates **(E)** derived from data presented in Figure 4. Linear prediction of bout probability in ulntA Air- and EtOH-exposed animals for the final four sessions of vapor exposure **(F)** and withdrawal **(G),** with beta estimates **(H)** derived from data presented in Figure 5. Linear prediction of bout probability in ContA EtOH-exposed and ulntA EtOH-exposed animals for the final four sessions of vapor exposure (I) and withdrawal **(J),** with beta estimates **(K)** derived from data presented in Figure 6. Error bars indicate ± 95% confidence intervals. Some error bars for beta estimates are small enough to be obscured by the mean. For models presented in this figure, the reference level for vapor group and schedule were Air-exposed and ContA. Real time or percent time were modeled as continuous predictor variables. Green or pink colored data points indicate statistically significant terms reflecting positive-moving or negative-moving beta estimates, respectively. Solid lines indicate ContA and dashed lines indicate ulntA.

**Supplementary Figure 3:**
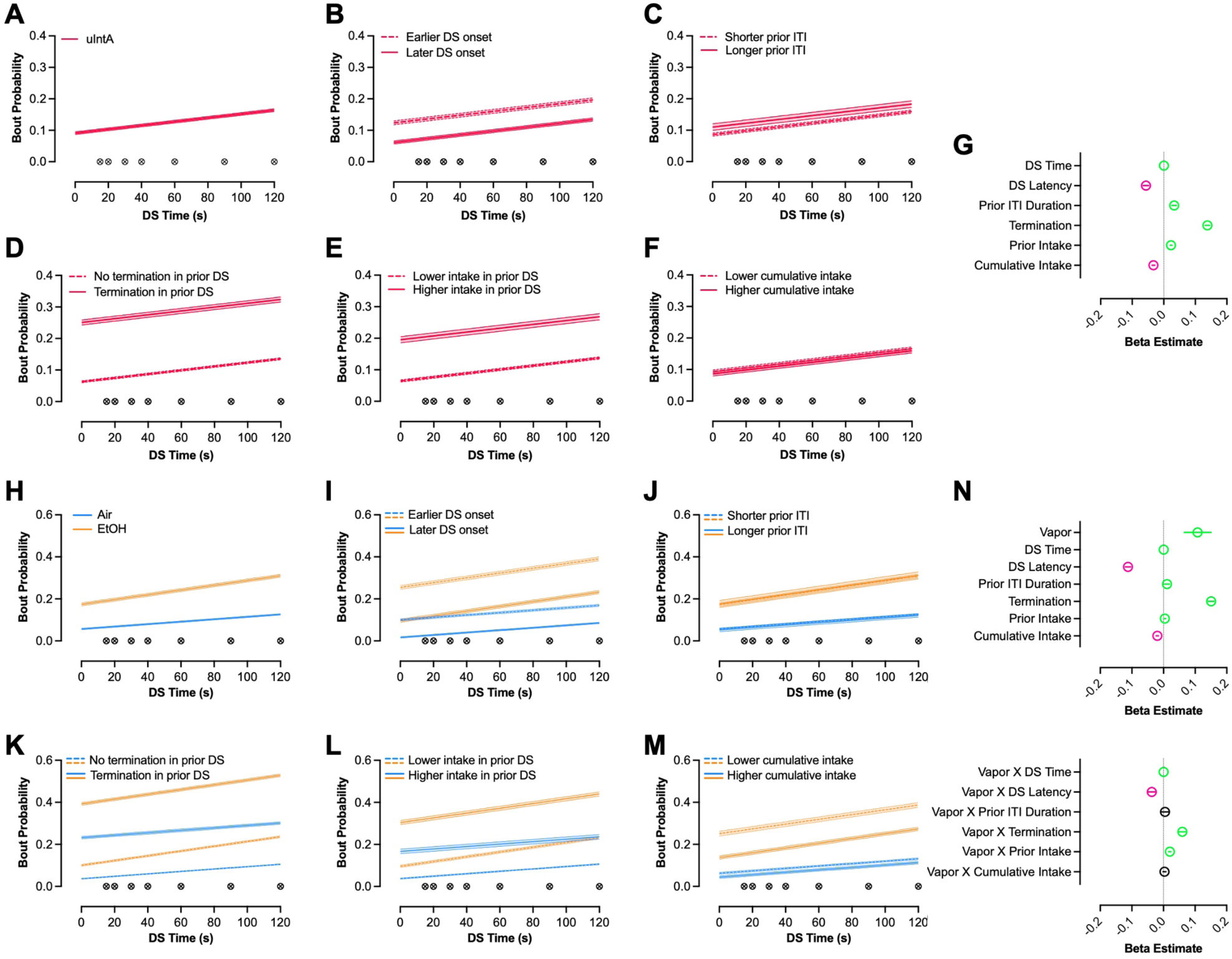
Linear predictions and beta estimates for variables affecting bout probability during DS presentations in the ulntA schedule. Linear prediction of bout probability in ulntA animals during the baseline timepoint across OS time **(A),** latency of OS onset **(B),** prior ITI duration **(C),** experience of a termination in the prior OS **(D),** ethanol intake in the prior OS **(E),** and cumulative intake during the session **(F),** as well as beta estimates **(G)** for each term. Linear prediction of bout probability in Air- and EtOH-exposed ulntA animals during the vapor and withdrawal timepoints across OS time **(H),** latency of OS onset **(1),** prior ITI duration **(J),** experience of a termination in the prior OS **(K),** ethanol intake in the prior OS **(L),** and cumulative intake during the session **(M),** as well as beta estimates **(N)** for each term *(top)* and their interaction with vapor group *(bottom).* Error bars indicate ± 95% confidence intervals. Some error bars for beta estimates are small enough to be obscured by the mean. For models presented in this figure, the reference level for vapor group was Air-exposed and the reference level for categorically defined OS-related parameters (i.e. prior ITI duration and experience of a prior termination) are set to the condition indicated by the dashed lines. All other OS-related parameters were modeled as continuous predictors. Green or pink colored data points indicate statistically significant terms reflecting positive-moving or negative-moving beta estimates, respectively. Inset symbols indicate unpredictable offset of the OS period (i.e. 15, 20, 30, 40, 60, 90, or 120 s).

**Supplementary Table 1:**
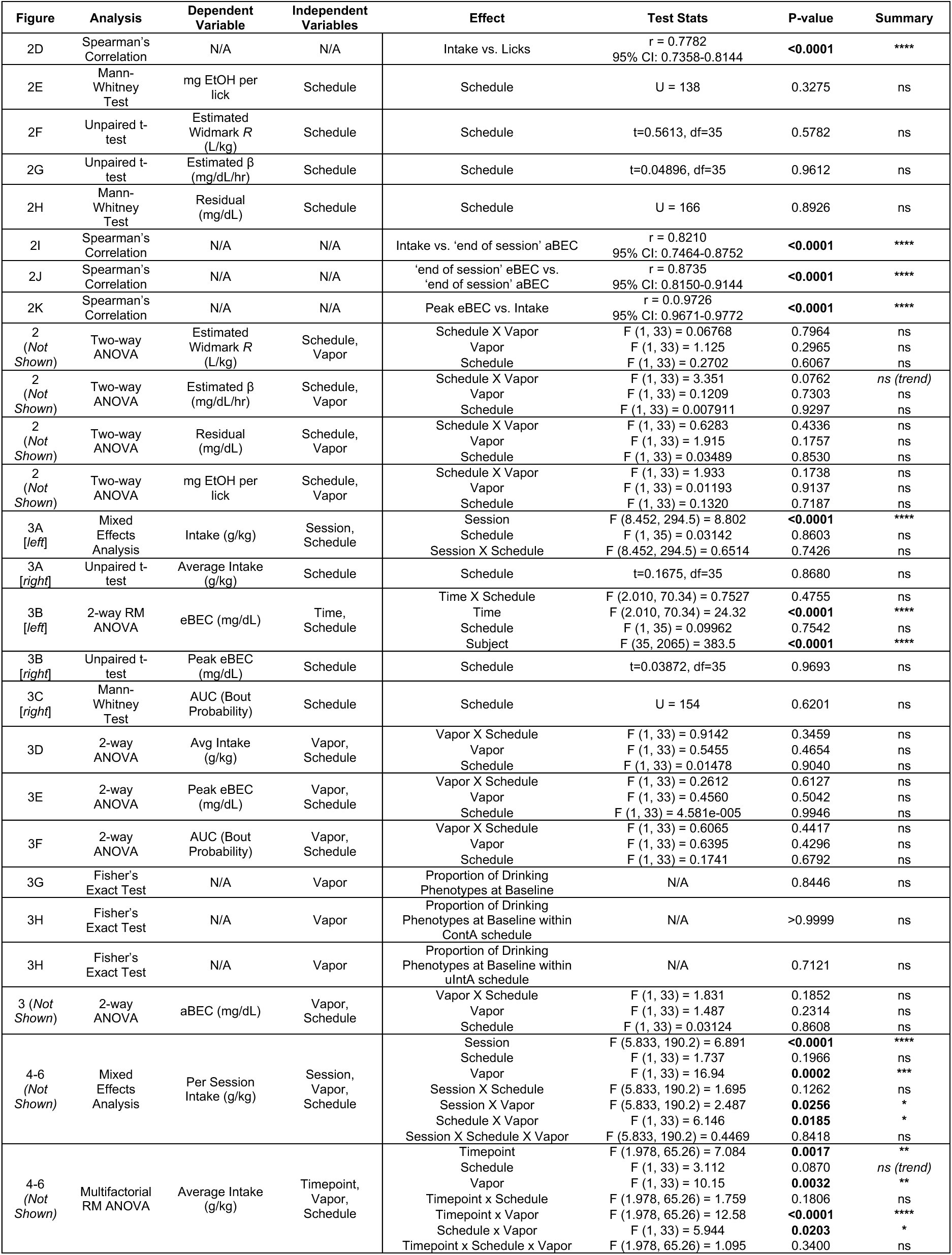
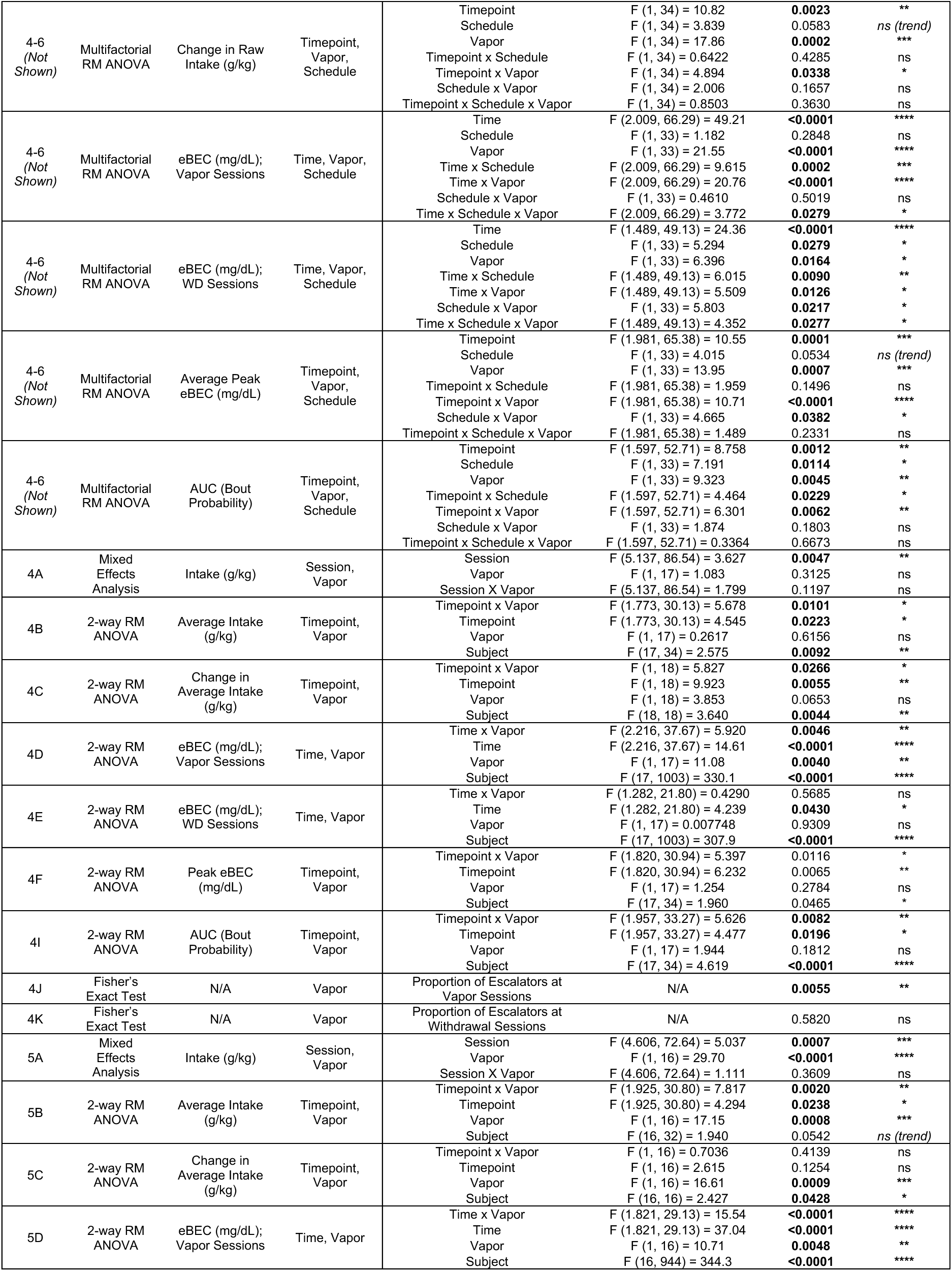
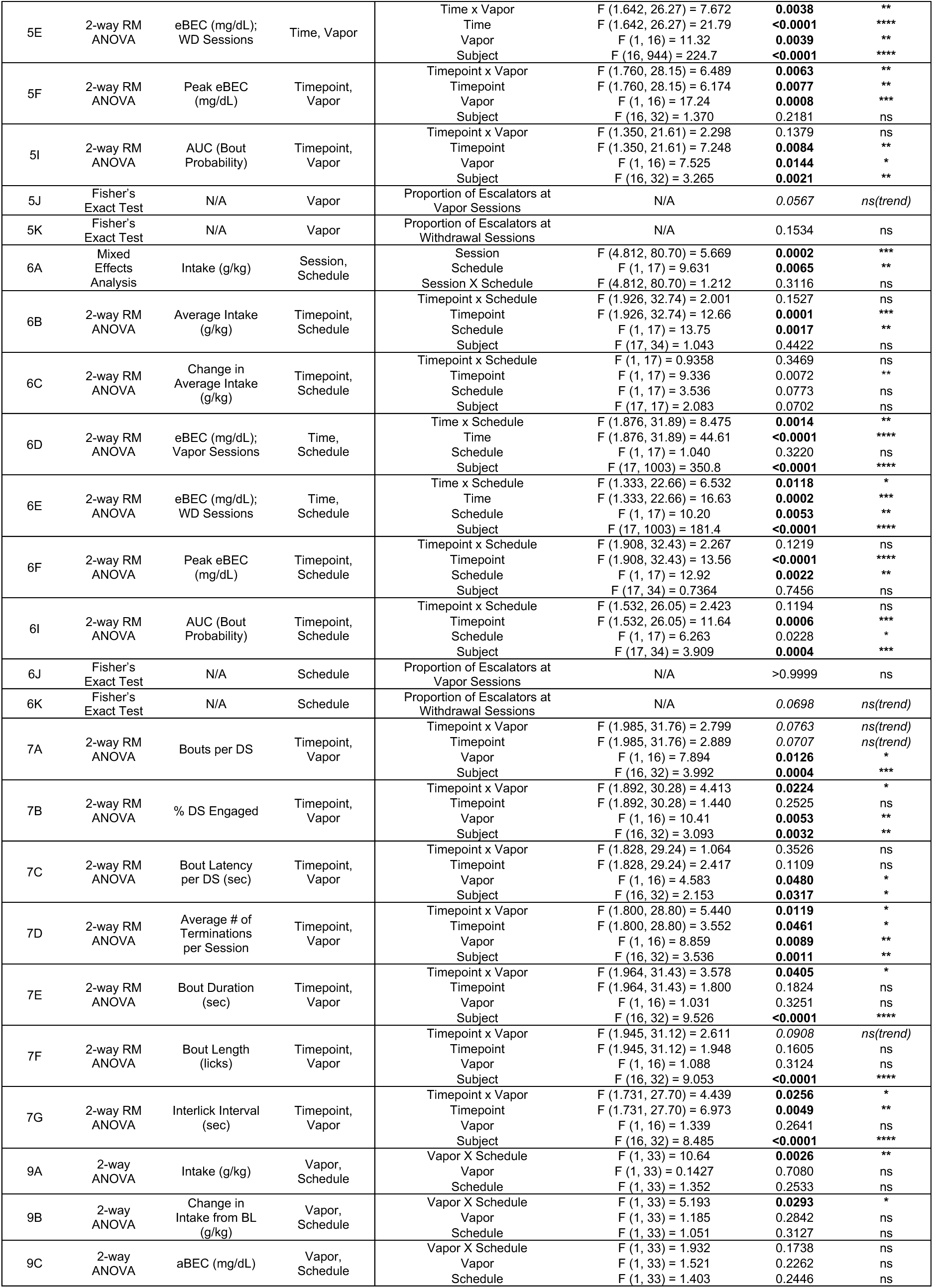
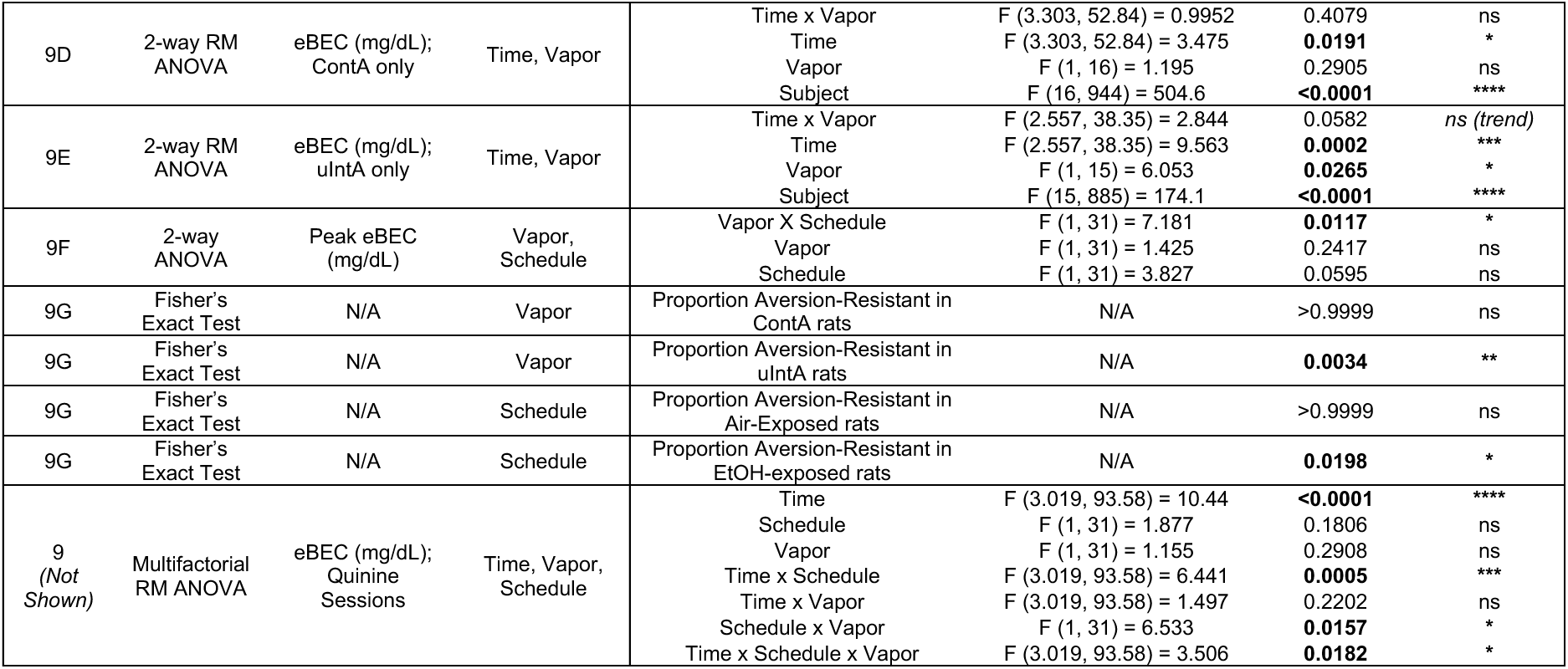
Results of all conventional statistical analyses.

**Supplementary Table 2:**
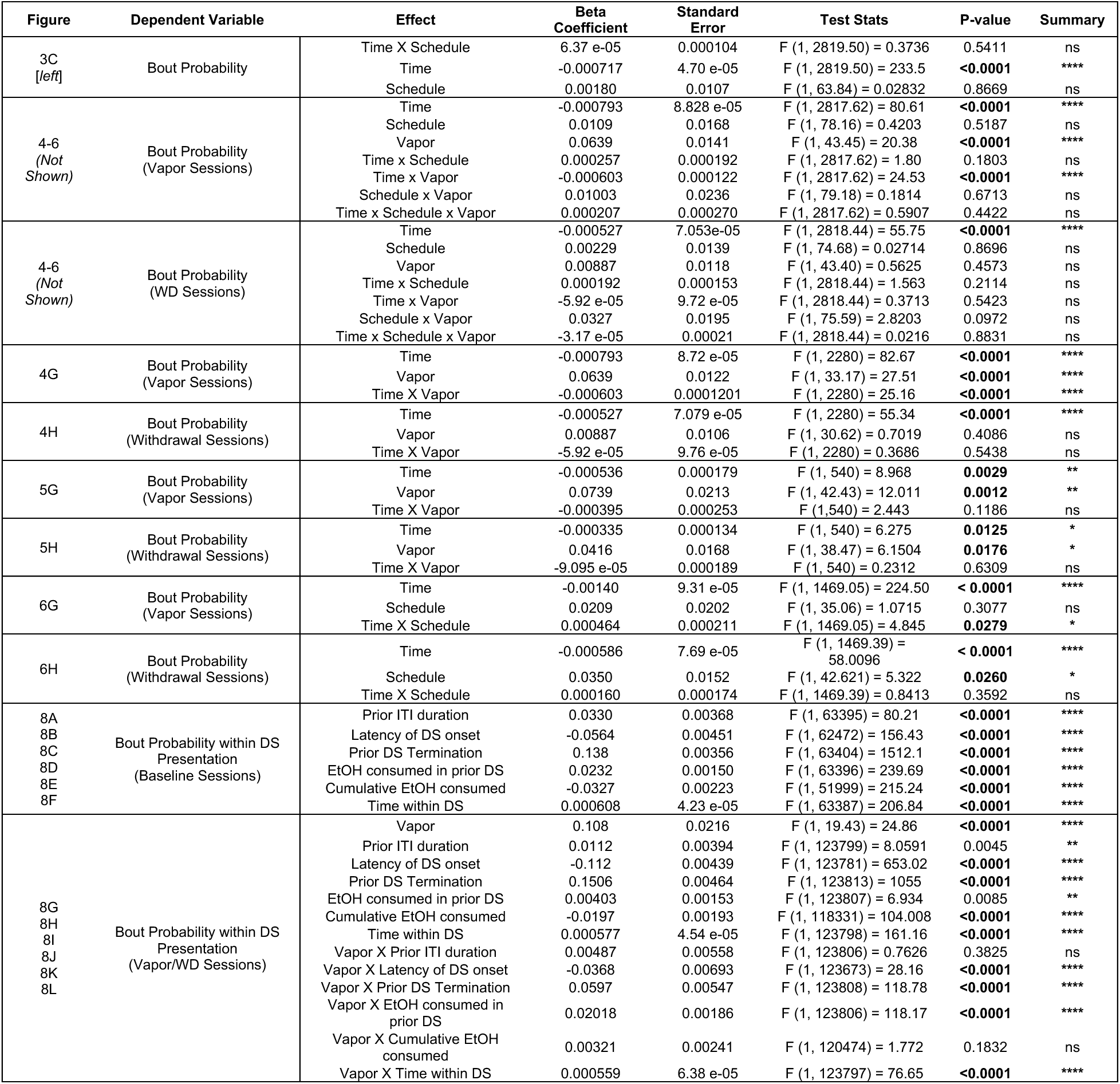
Results of linear mixed effect modeling.

